# *BRCA* mutation alters the stromal landscape in normal ovaries

**DOI:** 10.1101/2025.10.01.679597

**Authors:** Het T. Vaishnav, Alyssa Murray, Olivia R. Piccolo, Curtis W. McCloskey, Edward Yakubovich, Elizabeth Macdonald, Maryam Echaibi, Manon de Ladurantaye, Euihye Jung, Thomas J. Fazio, Dominique Trudel, Anne-Marie Mes-Masson, Ronny Drapkin, David A. Landry, Barbara C. Vanderhyden

**Author notes:** **Corresponding Author:** Barbara Vanderhyden. The authors contributed equally to this work (Het T. Vaishnav, Alyssa Murray).

## Abstract

For *BRCA* mutation (*BRCA+*) carriers, the risk of ovarian cancer can be as high as 59% compared with 1.4% for the general population. While the impact of *BRCA* mutations on epithelial cell transformation has been extensively studied, we hypothesize that these mutations cause structural changes that prematurely transform the ovary into a rich metastatic niche that supports the early onset of ovarian cancer. Analysis of collagen content and organization in human ovaries revealed increased coherence associated with fibrosis in premenopausal *BRCA*+ ovaries relative to those without a *BRCA* mutation. *Brca1* deficiency and haploinsufficiency in murine ovarian fibroblasts triggered the expression of hallmarks of senescence, including *Cdkn2a* (p16) and acidic β-galactosidase activity. *Brca1*-deficient fibroblasts also acquired an antigen-presenting myofibroblastic phenotype, characterized by expression of MHC-II molecules, α-SMA and extracellular matrix components, suggesting the capacity to modulate immune activity and drive structural changes resembling fibrosis. These results provide insight into the mechanisms contributing to accelerated ovarian aging in *BRCA+* carriers.

**Teaser:** *BRCA* mutation promotes fibroblast hyperactivity and senescence that changes the stromal architecture of the ovarian niche.

## Introduction

While cancer can develop at any age, it is considered an age-associated disease (*1*), with most cases being diagnosed after the age of 60 (*2*). The median age at diagnosis of sporadic ovarian cancer is 63 years, which is well after cessation of the ovarian functions required for fertility. However, for the 20-25% of ovarian cancer cases associated with germline mutations in the DNA repair genes *BRCA1* and *BRCA2*, the median age of onset decreases to 50-53 years (*3*). While the contribution of *BRCA1/2* mutations to driving malignant transformation of ovarian and fallopian tube epithelium has been well characterized (*4–6*), it is unclear whether this is the sole mechanism through which these mutations contribute to the early onset of ovarian cancer. A better understanding of the mechanisms driving the early onset of ovarian cancer among *BRCA1/2* mutation (*BRCA+*) carriers is necessary for the development of targeted prophylactic therapies that abrogate risk or delay onset.

One way to prevent ovarian cancer is to suppress the number of ovulations over a lifetime, most easily achieved by oral contraceptives. This becomes a difficult balance as *BRCA+* carriers have an increased risk of both breast and ovarian cancer. Oral contraceptives have been shown to decrease ovarian cancer risk in these patients; however, the risk of breast cancer increases due to the mitogenic effect of progestins in the breast (*7*). Since *BRCA* mutations already increase the risk of breast cancer up to 6 times that of non-mutation carriers, the use of oral contraceptives for ovarian cancer prevention is highly debated (*8*). As a result, clinicians and patients rely on enhanced screening or risk-reducing surgery options for early detection or prevention.

The ovary is one of the first organs to exhibit age-associated changes, characterized by a decline in fertility, followed by the cessation of ovulation and a reduction in estrogen production at natural menopause (*9*). Interestingly, the main non-hereditary risk factors for ovarian cancer are age, and the number of lifetime ovulations (*10–12*). We have previously discovered that ovarian fibrosis develops in an age-dependent manner in humans, evidenced by increased collagen linearization in postmenopausal ovaries relative to premenopausal ovaries (*13*). Tissue fibrosis, defined here as the accumulation and alignment of extracellular matrix (ECM) proteins, has been correlated with cancer development in the lung (*14*), breast (*15*, *16*), and liver (*17*). However, the mechanisms underlying the development of fibrosis in human ovaries have not been determined and its contribution to ovarian cancer is presently unknown.

Among the general population, the lifetime risk of developing ovarian cancer is approximately 1-2%; however, this risk is much higher for individuals who carry a germline mutation in *BRCA1* (39-63%) and/or *BRCA2* (11-27%) (*3*, *8*, *18*). Clinically, *BRCA+* carriers often exhibit a narrower reproductive window, characterized by reduced levels of anti-Müllerian hormone, fewer ovarian follicles and viable oocytes, and an earlier onset of menopause, which has been attributed to impaired DNA damage repair associated with these genes (*19–22*). We have previously suggested a link between ovarian fibrosis and reduced fertility (*23*) and others have shown that preventing or reversing fibrosis in mice can restore fertility (*24*).

While the majority of high-grade serous ovarian cancers (HGSC), the most common and lethal subtype, originate in the fallopian tube (*25*), tumors progress primarily after cells from serous tubal intraepithelial carcinomas (STICs) migrate to the ovary. It is now clear that the ovary significantly contributes to and may be required for the peritoneal dissemination of ovarian cancer, as ovariectomy can markedly attenuate peritoneal metastasis in fallopian tube-driven models of ovarian cancer (*26*, *27*). Further, up to 30% of cancers found in the ovary are metastatic from colon, stomach and breast (*12*, *28*). This demonstrates that the ovary is a pro-tumor niche, and the age-associated development of ovarian fibrosis that creates a cancer-supportive niche is a possible explanation.

Ovarian fibrosis has yet to be investigated in the context of high-risk women, specifically *BRCA+* carriers. This study aimed to explore the possibility that *BRCA1/2* mutations play a role in promoting a pre-metastatic niche in the ovary, beyond their established impact on epithelial cells-of-origin. Using two methods of histopathological imaging for collagen in an extensive cohort of well-curated human ovary samples, we show increased fibrotic content in cancer-free *BRCA+* carriers compared to non-carriers. Given that fibroblasts are the primary matrix remodelers and that persistent myofibroblast activation is strongly associated with the development of pathological fibrosis (*29*, *30*), we investigated the impact of *Brca1* deficiency on myofibroblast activation. Using RNA-sequencing, we show that the loss of *Brca1* in fibroblasts induces a unique senescent, myofibroblastic, antigen-presenting cancer-associated fibroblast (CAF)-like phenotype, which may be associated with continual matrix deposition and disrupted organization. These results, validated in fibroblasts with heterozygous loss of *Brca1* reveal a mechanistic underpinning for the creation of a primed ovarian niche that may help to drive early cancer onset in *BRCA+* carriers.

## Results

### *BRCA1/2* mutation supports the development of early onset ovarian fibrosis

To assess collagen content and organization, ovarian cortex sections from cancer-free patients were stained with picrosirius red staining and imaged under polarized light (PSR-POL). *BRCA* mutation was used as a comparator within premenopausal patients who ranged in age from 35-49 (N = 48; Supplementary Table 1 shows age and BRCA status). When collagen content was assessed, we observed variation within both groups, however collagen was present in at least 30% of the stromal area in 75% of all samples. While no difference in the abundance of collagen was found (Fig. 1A), the coherency of the collagen fibers was significantly more aligned, indicative of fibrosis, in the group with a *BRCA* mutation compared with those without (Fig. 1B; p = 0.0076). Patients were stratified by the BRCA protein mutated (BRCA1 or BRCA2); however, no statistically meaningful differences were observed between the cohorts. Representative PSR-POL images for low and high coherency and abundance values are presented in Figs. 1C-F. Linear regression showed poor correlation between coherency and abundance across samples (R^2^ = 0.25).

**Fig 1.**
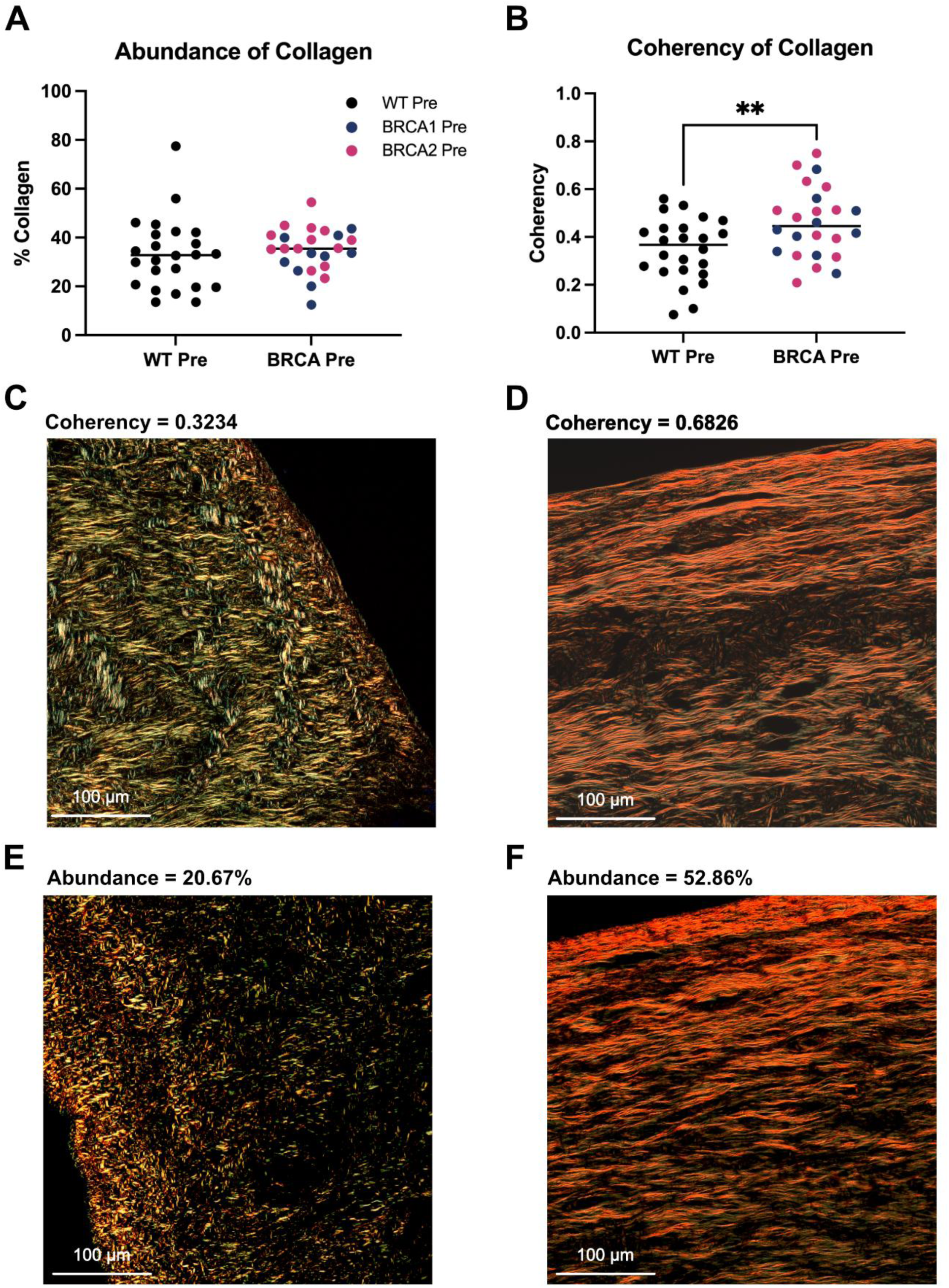
PSR-POL reveals *BRCA* mutation is associated with increased ovarian fibrosis in cancer-free patients. **(A)** Total collagen within selected region of interest (ROI) as determined by HUE program on Fiji. **(B)** Coherency of collagen fibers within selected ROI as determined by the OrientationJ Dominant Direction program on Fiji. Representative images of low and high coherency (**C**, **D**) and abundance (**E**, **F**) for ovarian cortex regions assessed by PSR-POL are shown. Analysis was completed on ovaries from cancer-free premenopausal patients (aged 35-49) with and without *BRCA* mutation. Patients with a *BRCA1* mutation are identified in blue while patients with a *BRCA2* mutation are identified in pink. Significance was determined by unpaired t-test and shown by ** p < 0.01 (*n* = 24).

### Single-fiber analysis of collagen within the ovarian cortex with the digital pathology platform FibroNest

We further characterized the collagen structure through single-fiber, high resolution analysis of collagen within the ovarian cortex using the digital pathology platform, FibroNest. We analyzed five ovaries per condition, including premenopausal without known mutation (WT Pre), premenopausal with a *BRCA1/2* mutation (BRCA Pre) and postmenopausal without a BRCA mutation (WT Post). The ovaries were selected to encompass a representative range of coherency values within each group (at, two above, and two below median) from the PSR-POL experiment.

The full analysis included scores for various aspects of collagen content (the density, reticulation and quantity of total vs. fine vs. assembled collagen), morphometry (fiber length, thickness, area, perimeter) and architecture (organization, compactness and patterns). Of the 177 quantitative fibrosis traits (qFTs) calculated by FibroNest, 50 were significant and unique to age (WT Post compared with WT Pre), 7 were significant and unique to BRCA (WT Pre vs BRCA Pre) and 6 were significant and common to both the age and BRCA groups. All significant qFTs (p < 0.05) are displayed in a phenotypic heat map (Fig. 2). A representative heat map identifying the directionality of the qFT changes as compared to WT Pre can be found in Supplementary Fig. 1. Of the 50 qFTs significant to age, the amount of assembled collagen increased in WT Post compared with WT Pre. Traits associated with fine collagen fibers decreased in their scores while qFTs associated with assembled collagen showed increases. This led to variation in the directionality of scores associated with all collagen fiber qFTs. There were several properties found in BRCA Pre ovaries that were not found in the WT Pre vs WT Post analysis. These distinct properties included changes to the tortuosity and orientation of the collagen fibers. This occurred within all collagen but was more pronounced in fine collagen fibers. Common to both age and BRCA groups were changes to the eccentricity of the collagen fibers, especially amongst fine fibers. Interestingly, not all of the BRCA Pre ovaries displayed the same phenotype with patients 32 and 39 appearing most similar to WT Post while patients 25, 37 and 44 share more characteristics with WT Pre.

**Fig 2.**
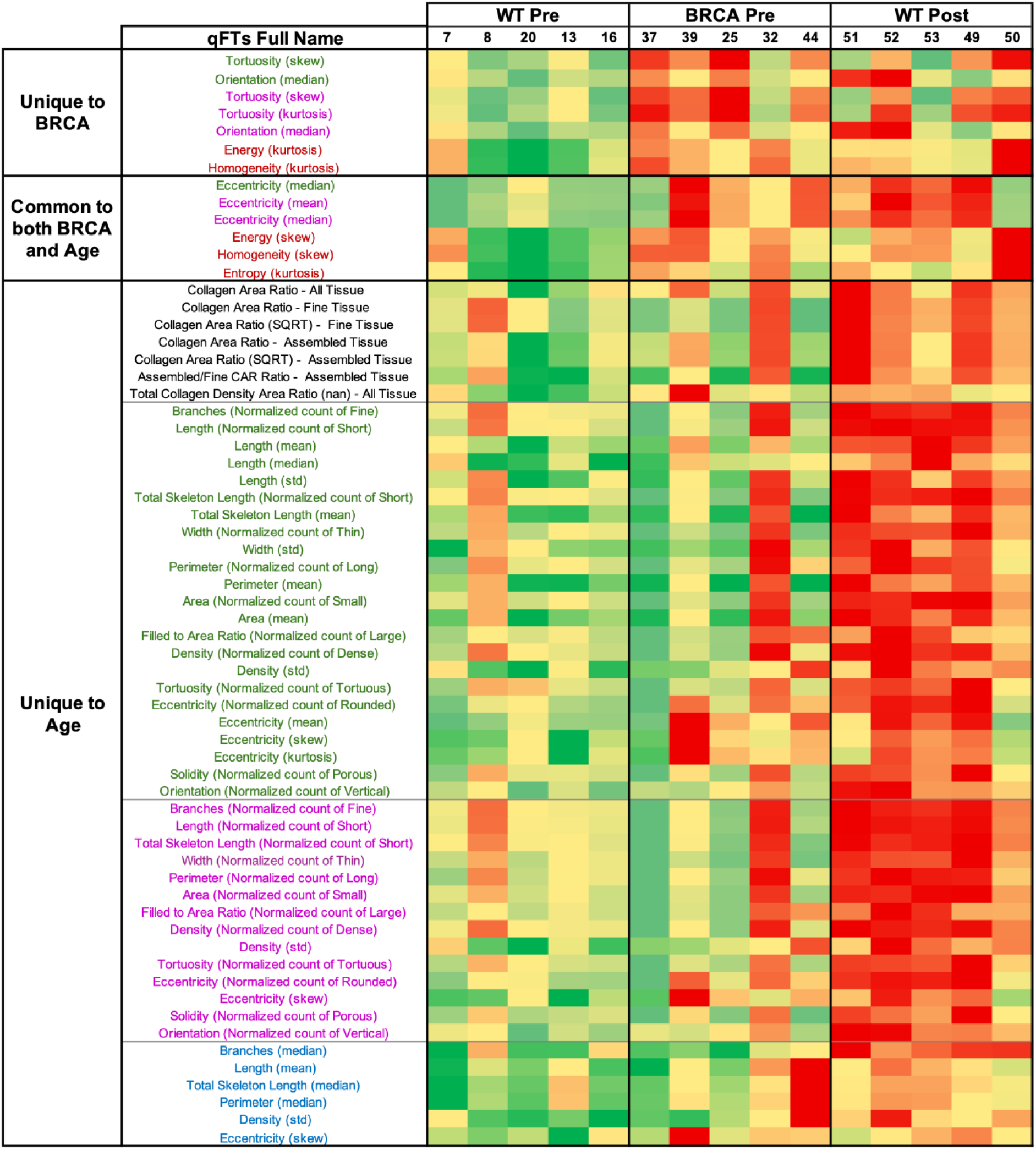
Single fiber analysis using the FibroNest digital pathology platform identifies differences in ovarian cortex stratified by age and *BRCA* mutation. Phenotypic heatmap of fibrosis traits for each sample as identified by the FibroNest platform. Only significant quantitative fibrosis traits (qFTs; p < 0.05) are presented and are segregated based on being unique to BRCA, common to both BRCA and age and unique to age. Numbers within each group identify patients as listed in Supplemental Table 1. Red indicates difference in trait value from Pre WT while green indicates similarity in trait value to Pre WT. qFT colour descriptions are as follows: black – collagen content; green – collagen morphometry among all fibers; pink – collagen morphometry among fine fibers; blue – collagen morphometry among assembled fibers; red – fibrosis architecture. Significance was determined by Welch’s t test (*n* = 5).

Representative images of the FibroNest analysis for each group (WT Pre, BRCA Pre and WT Post) are shown in Fig. 3A. The fiber density, collagen architecture and size of collagen patches all appeared significantly different in the WT Post group compared with the WT Pre. Within the BRCA Pre group, changes to these parameters were also evident compared with WT Pre, however, were not as pronounced as in the WT Post group.

**Fig 3.**
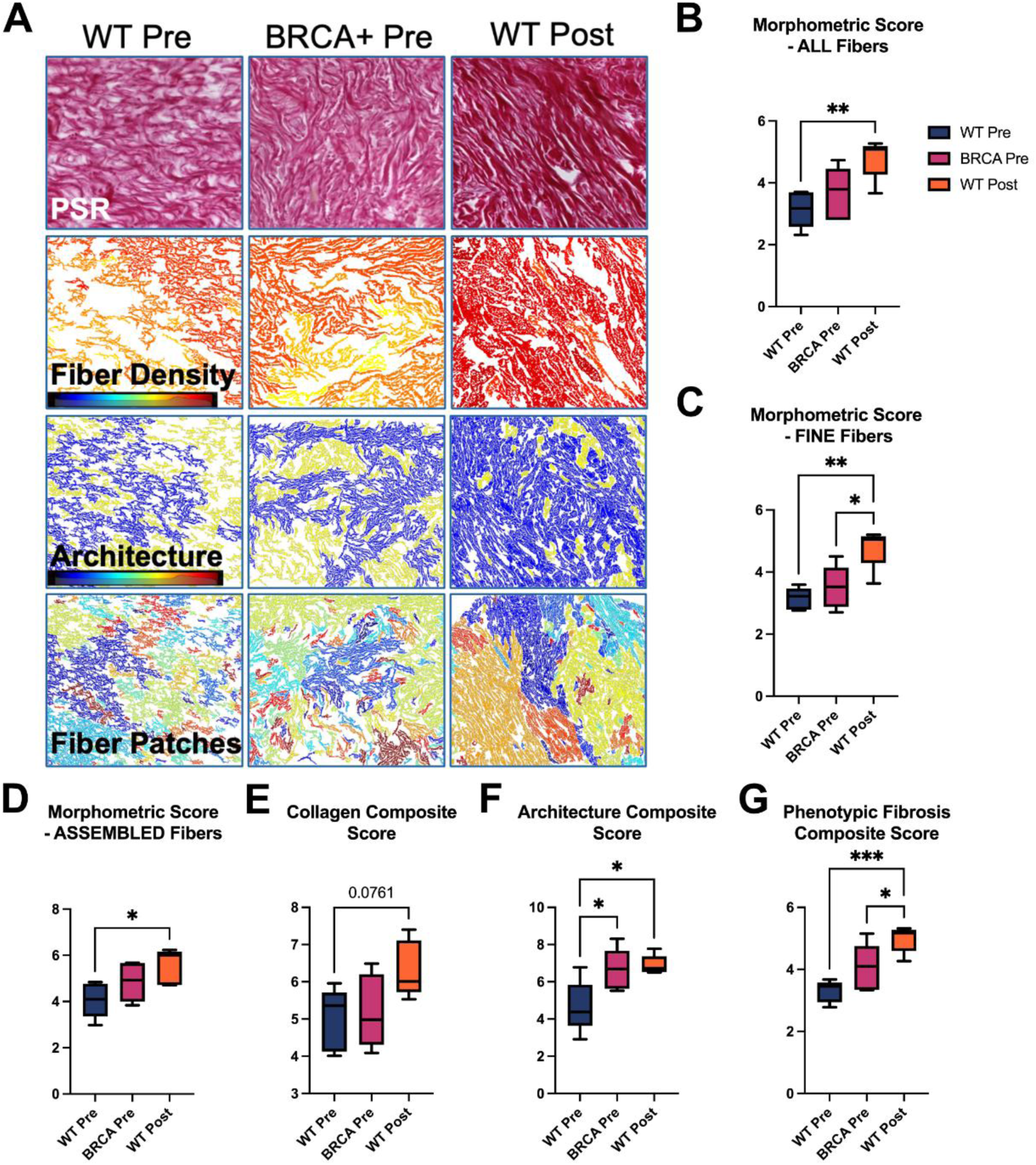
Fibrosis scores calculated by FibroNest confirm *BRCA* mutation alters the collagen architecture. **(A)** Representative images of the FibroNest analysis for each of the groups (WT Pre, BRCA Pre and WT Post) are shown for brightfield PSR imaging, fiber density, collagen architecture and collagen fiber patches. Principal quantitative fibrosis traits were combined into normalized composite fibrosis scores using the FibroNest digital AI platform. Morphometry was separated by collagen type: all fibers **(B)**, fine fibers **(C)**, or assembled fibers **(D)**. Collagen content is represented by collagen composite score **(E)**, collagen architecture is reflected by architecture composite score **(F)**, and overall fibrosis is represented by the phenotypic fibrosis composite score **(G)**. Analysis was completed on ovaries from premenopausal patients with (BRCA Pre) and without (WT Pre) a *BRCA* mutation as well as postmenopausal patients without a *BRCA* mutation (*n* = 5). One way ANOVA with Tukey’s post-hoc test was used to determine significance between groups. * p < 0.05, ** p < 0.01, *** p < 0.001.

The qFTs were also used to calculate various fibrosis scores based on the severity of fibrosis within the sample. As described above, collagen morphometry was assessed in three parts. There was a significant difference with age amongst all collagen phenotypes (total; Fig. 3B, fine; Fig. 3C and assembled; Fig. 3D), however, there was no difference with *BRCA* mutation. Collagen composite score assessed the collagen content within the sample and there was no significant difference between groups and large variability among samples (Fig. 3E), although there was a strong trend (p=0.0761) for this score to be higher in WT Post samples. This matches what was found using PSR-POL and strengthens the notion that *BRCA* mutation does not affect total collagen abundance. The collagen architecture composite score, however, showed significant differences in both age and *BRCA* mutation groups compared to WT Pre (Fig. 3F). This is in concordance with increased coherence shown in Fig. 1B and is represented visually in Supplementary Fig. 2. Lastly, the phenotypic fibrosis composite score showed a significant difference between WT Pre and Post with BRCA Pre being intermediate, exhibiting a larger degree of variation (Fig. 3G). The entire FibroNest analysis mirrors what we have reported previously using different methods (14); that human ovaries develop age-associated fibrosis. The results also indicate that premenopausal ovaries with *BRCA* mutation develop some, but not all of the features that develop with age, but the key feature that is shared by *BRCA* mutation and age is collagen orientation (coherence) which consistently reflects fibrosis.

### Adenoviral cre-mediated recombination creates *Brca1* KO MOFs

To explore the consequences of BRCA dysfunction in ovarian fibroblasts, a likely source for the disrupted ECM structure in human *BRCA* mutant ovaries, we altered *Brca1* expression in isolated murine ovarian fibroblasts (MOFs). *Brca1*^LoxP/LoxP^ conditional knockout mice, bearing LoxP sites in introns 4 and 13 of the *Brca1* gene, were used as a source for MOFs. To validate the system, enhanced yellow fluorescent protein (EYFP) expression from the Rosa26 locus, was tracked as a reporter of MOFs in which cre-mediated recombination of *Brca1* had occurred. EYFP expression was detected starting 2 days post-transduction. By 14 days, stable EYFP expression was confirmed in MOFs transduced by AdCre (*Brca1*-KO), but not in MOFs transduced by AdEmpty (*Brca1*-WT) (Fig. 4A). Genomic DNA and mRNA extracted from *Brca1*-KO and *Brca1*-WT MOFs were used to assess genomic recombination and expression of *Brca1*. Amplicon products for both recombined and WT *Brca1* were detected in gDNA from AdCre-transduced MOFs, with the latter to a much lesser extent (Fig. 4B). Only WT *Brca1* was detected in gDNA from AdEmpty-transduced MOFs (Fig. 4B). Quantitative PCR (qPCR) analysis revealed a significant decrease in the mRNA expression of *Brca1* after AdCre transduction, as compared to AdEmpty-transduced MOFs (Fig. 4C), suggesting successful *Brca1* KO and loss of expression in most cells. The KO was further confirmed by RNA-sequencing, which revealed significantly reduced mRNA expression in *Brca1*-KO MOFs (Fig. 4D).

**Fig 4:**
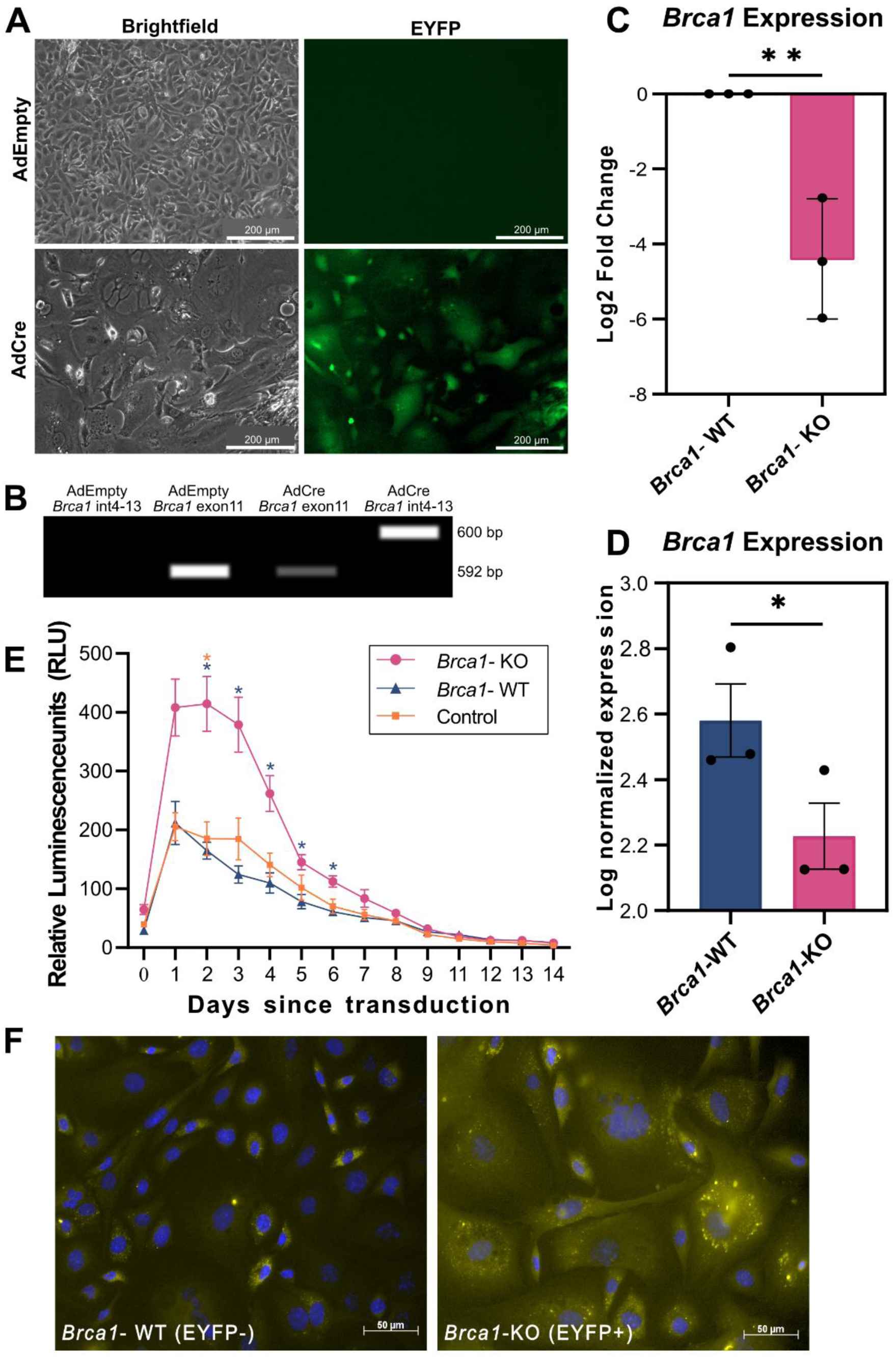
Deletion of *Brca1* induces apoptosis or a senescence-like phenotype in MOFs. MOFs (passage=1) were transduced with either AdCre or AdEmpty as control. EYFP expression (used to track *Brca1* deletion) and cell death by apoptosis were monitored daily for 14 days, after which cells were collected for gDNA and RNA extraction to confirm genomic recombination and mRNA expression of *Brca1* as well as RNA-sequencing. Experiment was performed with three biological replicates (n=3). **(A)** 14 days post-transduction AdCre MOFs strongly express EYFP and display a unique flattened morphology, while AdEmpty MOFs retain their normal morphology and do not express EYFP. **(B)** PCR using gDNA from AdCre MOFs produced two amplicons, a 592bp product from wild-type (wt) *Brca1* (primer set targets exon 11), and a 600bp product from cre-mediated recombination of *Brca1* (primer set targets intron4-13 junction), while AdEmpty MOFs retain the 592bp wt product only. **(C)** Quantitative PCR (qPCR) revealed significantly decreased mRNA expression of *Brca1* in AdCre MOFs as compared to AdEmpty MOFs. Raw data normalized using housekeeping genes *Act-β*, *Gapdh* and *Hrpt1*, analyzed using the relative quantification (2-ΔΔCt) method and presented as relative fold change (Log2 fold change; Log2FC). (**D**) RNA-sequencing confirms decreased mRNA expression of *Brca1* in AdCre MOFs, relative to AdEmpty MOFs. Relative abundance of *Brca1* mRNA is shown as log normalized expression between *Brca1*-KO and *Brca1*-WT MOFs. **(E)** Cell death was measured using RealTime-Glo Apoptosis Assay kit every 24h, with time point 0 days representing 4h post-transduction. At all time points, apoptosis was recorded to be highest in *Brca1*-KO MOFs, compared to *Brca1*-WT or control (non-transduced) MOFs. **(F)** Immunofluorescence reveals development of senescence-like cellular morphology, characterized by increased cell size due to an enlarged morphology, and abnormal nuclei in *Brca1*-KO MOFs, while *Brca1*-WT MOFs retained their wild-type fibroblast morphology and do not express EYFP. *Brca1*-WT MOFs appear to exhibit a little auto-fluorescence when excited using a wavelength of 488nm. Nuclei were stained using Hoechst. Scale bar = 50 µm. Biotek Cytation 5 imaging reader was used to visualize EYFP and capture images at 10x (**A**) and to measure luminescence (**E**). Axioskop 2 MOT fluorescence microscope was used to capture images in (**F**). Error bars indicate standard error of mean (SEM). Unpaired student’s t-test (**C** and **D**) and two-way ANOVA (**E**) were applied using GraphPad Prism and R (**D**); *= p < 0.05 and **= p < 0.01.

### Sudden loss of *Brca1* leads to cell death via apoptosis, while surviving cells show enlarged and flat morphology

Using a luminescence-based apoptosis assay, a significant increase in apoptosis was observed in *Brca1*-KO MOFs compared to *Brca1*-WT MOFs from 2 to 6 days post-transduction, after which no significant differences were observed between the two groups, suggesting that the sudden loss of BRCA1 function induces apoptosis in MOFs (Fig. 4E). Cell death by apoptosis peaked 2 days post-transduction in *Brca1*- KO, gradually decreasing with time and reached a plateau at day 14 post-transduction, suggesting recovery from viral transduction.

Surviving *Brca1*-KO MOFs adopted an enlarged and flattened morphology, compared to *Brca1*-WT MOFs that retained their normal morphology (Fig. 4F). This observation suggests that cells that were able to withstand the sudden loss of *Brca1* survived but adopted a morphology that is characteristic of senescent cells.

### *Brca1* KO increases the expression of genes associated with genome organization and immune regulation pathways in MOFs

To assess transcriptomic changes associated with the loss of *Brca1* in MOFs, we conducted RNA-sequencing. Gene Set Enrichment Analysis (GSEA) of differential genes expressed in *Brca1* KO MOFs revealed significant enrichment of processes and pathways involving genomic reorganization, assembly and disassembly of the organizational units of the genome, as well as processes mediating immune regulation and a significant decrease in the processing of non-coding mRNA (Fig. 5A). The upregulation of pathways involved in genome reorganization and chromatin assembly/disassembly, accompanied by the loss of BRCA1 function, aligns with its well-established role in DNA repair and the maintenance of genomic integrity.

**Fig 5:**
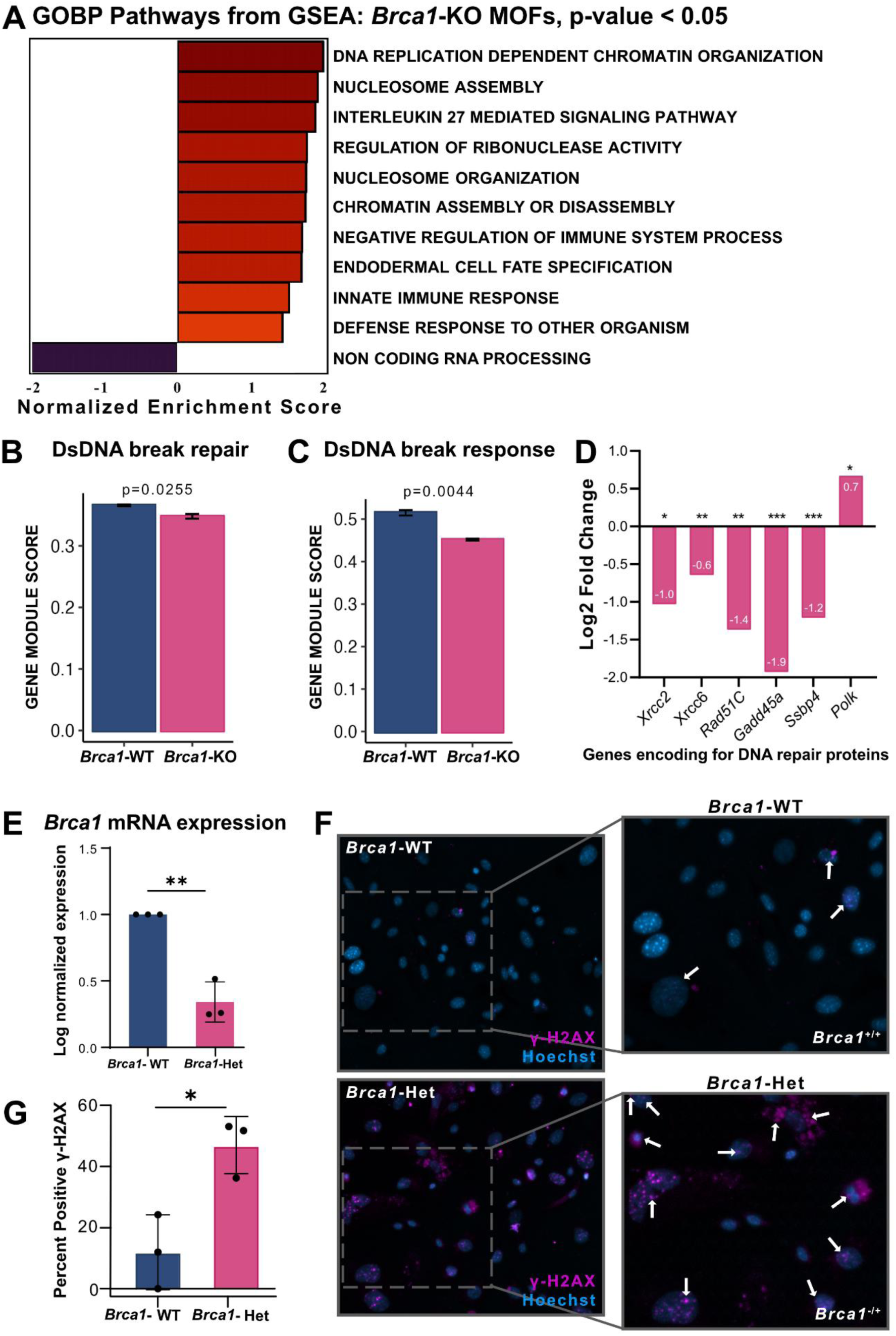
Loss of *Brca1* in MOFs impairs DNA damage repair pathways, leading to the accumulation of dsDNA damage. *Brca1*^LoxP/LoxP^ MOFs were transduced with either AdCre (*Brca1*-KO) or AdEmpty (*Brca1*-WT) and RNA was extracted 14 days post-transduction for RNA-seq analysis. Experiment was performed in triplicate (n=3). **(A)** Gene Ontology Biological Process (GOBP) pathways from GSEA. The loss of *Brca1* in MOFs decreased the gene module score for genes involved in (**B**) dsDNA break repair and (**C**) Cellular response to dsDNA damage, as compared to *Brca1*-WT MOFs. (**D**) The mRNA expression of various DNA repair proteins was significantly decreased with loss of *Brca1*, presented as log2FC in *Brca1*-KO MOFs, compared to *Brca1*-WT MOFs. (**E**) Quantitative PCR (qPCR) confirms reduced expression of *Brca1* in *Brca1*-Het MOFs, as compared to *Brca1*-WT MOFs. Raw data normalized using housekeeping genes *Act-β*, and *Ppia*, analyzed using the relative quantification (2-ΔΔCt) method and presented as log normalized expression. (**F**) *Brca1*-WT and *Brca1*-Het (*Brca1*^-/+^) MOFs were fixed using 4% paraformaldehyde for immunofluorescence staining of γ-H2AX 14 days post-transduction. γ-H2AX positive foci are visualized in pink and identified by white arrows and are largely absent in *Brca1*-WT MOFs. Nuclei were stained using Hoechst. Images captured using Axioskop 2 MOT fluorescence microscope, and percentage of cells positive for γ-H2AX was quantified (**G**). Error bars indicate SEM, Unpaired student’s t-test was performed using R (**A-D**) and GraphPad Prism (**E**). All gene sets were acquired from MSigDb and scored using Singscore in R project (**B**, **C**). *= p < 0.05, **= p < 0.01, ***= p < 0.001.

### Heterozygous loss of *Brca1* leads to accumulation of DNA damage in MOFs

The role of BRCA1 in maintaining genomic integrity and in orchestrating DNA repair pathways is well established, with *Brca1* mutant cells exhibiting impaired double stranded DNA (dsDNA) repair (*22*). To determine the consequences of BRCA1 loss on dsDNA repair processes in MOFs, RNA-seq data were analyzed using established gene sets from MSigDb associated with various DNA repair processes. These were scored using Singscore to generate a gene module score, which, for a given gene set is a rank-based average of the expression of all genes within that gene set (*31*, *32*). The loss of *Brca1* significantly impaired the gene expression (gene module score) of proteins required for dsDNA break repair and those required to evoke a cellular response to dsDNA damage (Fig. 5B, C), thus confirming the role of BRCA1 in MOFs as a key mediator in the assembly and function of DNA repair complexes. Decreased expression of genes encoding various DNA damage repair proteins, such as *Xrcc2*, *Xrcc6*, *Rad51c*, *Gadd45a* and *ssbp4* (Fig. 5D), suggested that BRCA1 is not only an integral component for DNA repair by homologous recombination but may also be responsible for the transcriptional regulation of DNA repair proteins required for other DNA damage repair pathways, such as non-homologous end joining and base excision repair. There was also an increase in the expression of *Polk*, which encodes for DNA polymerase κ, a specialized error-prone polymerase that is implicated in the repair of interstrand cross links, co-localizes with DNA repair proteins at sites of DNA damage, and is equipped to bypass dsDNA damage to continue strand extension (*33–35*). This suggests that the loss of *Brca1* likely leads to the accumulation of irreparable DNA damage.

Since the majority of human carriers are heterozygous for *BRCA1* mutations and have one functional copy (at least prior to tumorigenesis) (*36*), we assessed whether heterozygous loss of *Brca1* also leads to accumulation of DNA damage. Using qPCR, we confirmed the reduced expression of *Brca1* in heterozygous MOFs (*Brca1*^-/+^; *Brca1*-Het MOFs, Fig. 5E). We performed immunofluorescence staining for phosphorylated histone variant H2Ax (γ-H2AX), a marker of dsDNA damage, in MOFs heterozygous for the loss of *Brca1*. Significantly more γ-H2AX positive foci were observed in the nuclei and cytoplasm of *Brca1*- Het MOFs and were largely absent in *Brca1*-WT MOFs (Fig. 5 F,G). This confirms that heterozygous loss of BRCA1 in MOFs results in the accumulation of dsDNA damage, a well-known trigger for cellular stress and cell cycle arrest (*37*).

### Cell cycle arrest in *Brca1*-KO MOFs

Accumulation of DNA damage can lead to genomic instability and trigger cellular stress responses, such as cell cycle arrest to limit the propagation of compromised genomes that can drive malignant transformation of cells (*38*). *Brca1*-KO MOFs showed increased expression of genes (gene module score) involved in the inhibition of DNA replication mediated via phosphorylated retinoblastoma protein (pRb)-E2f complex (Fig. 6A). This complex is a key component of the G1/S checkpoint pathway that quenches E2F, a key transcription factor responsible for initiating DNA replication, suggesting inhibition of cell cycle progression in *Brca1*-KO MOFs. This was consistent with decreased expression of *Pold1*, *Pold2*, and *Pold4*, all of which encode for subunits of replicative DNA polymerase δ, as well as *E2f5*, a member of the E2f family of cell cycle-promoting transcription factors (Fig. 6B). Additionally, we noted increased expression of genes associated with cell cycle inhibition, such as *Rb1*, which encodes for Rb, an inhibitor of the E2f family of transcription factors, as well as several cyclin-dependent kinase inhibitors, specifically *Cdkn1a*, *Cdkn2a*, *Cdkn1c*, and *Cdkn2b*, encoding for p21, p16, p57 and p15, respectively (Fig. 6B). This concurrent increase in the expression of cell cycle inhibitors and decreased expression of cell cycle promoters reflect replicative arrest in *Brca1*-KO MOFs.

**Fig 6:**
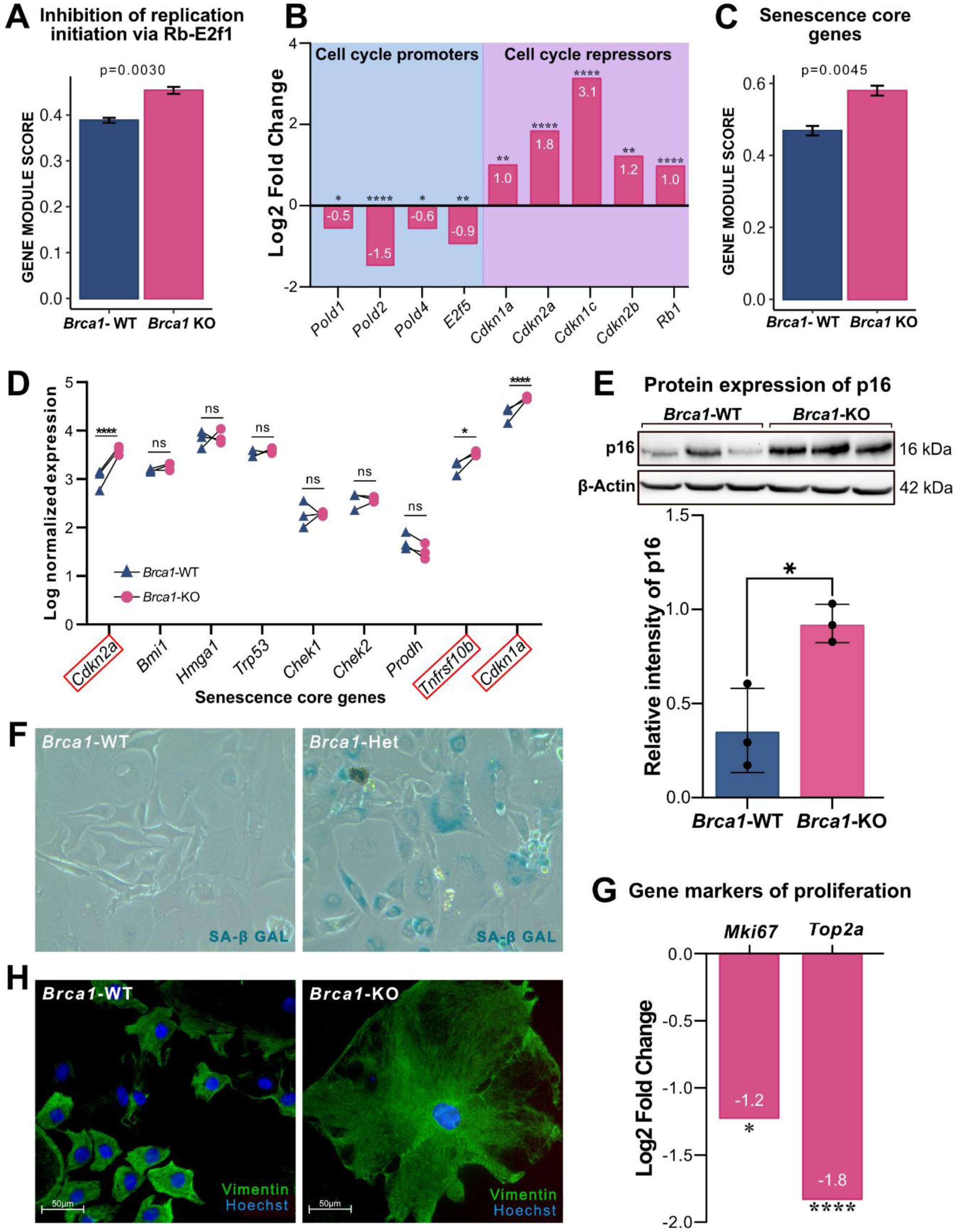
Loss of *Brca1* in MOFs results in the induction of cellular senescence. *Brca1*-KO and *Brca1*-WT MOFs were harvested 14 days post-transduction to extract RNA for RNA-sequencing (n=3) and 7 days post-transduction to extract protein for western blotting (n=3). (**A**) Gene module score for genes involved in “Inhibition of replication initiation via Rb-E2f1” was significantly upregulated by *Brca1*-KO MOFs, compared to *Brca1*-WT MOFs. (**B**) *Brca1*-KO MOFs show reduced expression of genes involved in DNA replication and cell cycle progression (shown in green), and increased expression of genes required for cell cycle checkpoint inhibition (shown in red), which are presented as Log2FC. (**C**) Gene module score for “Senescence core genes” is significantly upregulated in *Brca1*-KO MOFs, compared to *Brca1*-WT MOFs. (**D**) The expression of three core senescence initiator genes - *Cdkn2a*, *Tnfrsf10b*, and *Cdkn1a* - was significantly upregulated with loss of *Brca1*, presented as log normalized expression between *Brca1*-WT and *Brca1*-KO MOFs. (**E**) *Brca1*-KO MOFs show significantly increased expression of p16 compared to *Brca1*-WT MOFs 7 days after transduction. P16 expression was normalized using β-Actin and is presented as relative intensity. (**F**) *Brca1*-Het and *Brca1*-WT MOFs were stained for senescence-associated β-galactosidase activity 14 days post-transduction. (**G**) *Brca1*-KO MOFs show significantly decreased expression of genes encoding for markers of proliferation, Ki-67 and DNA topoisomerase II-α, presented as Log2FC. (**H**) Immunofluorescence staining of vimentin, an intermediate cytoskeletal filament highlights enlarged, spread-out cell bodies of *Brca1*-KO MOFs (right), whereas *Brca1*-WT MOFs retain their normal morphology (left). Nuclei were stained using Hoechst. Scale bar, 50 µm. Error bars indicate SEM unpaired student’s t-test was performed using R (**A-D, G**) and using GraphPad Prism for (**E**). *= p < 0.05, **= p < 0.01, ***= p < 0.001, ****= p < 0.0001.

### Loss of *Brca1* is associated with the induction of cellular senescence in MOFs

Given that the loss of *Brca1* caused MOFs to transition to an enlarged and flat cellular morphology alongside increased expression of cell cycle inhibitors, we assessed whether the loss of *Brca1* resulted in onset of cellular senescence. The gene module score of senescence core genes was significantly upregulated in *Brca1*-KO MOFs (Fig. 6C), particularly due to upregulation of *Cdkn2a* (p16; log2FC=1.83), *Cdkn1a* (p21; log2FC=1.00), and *Tnfrsf10b* (TRAIL2; log2FC=0.88), identifying them as core senescence initiators associated with *Brca1* loss in MOFs (Fig. 6D). Western blots confirmed the increased expression of p16, which was detected as early as 7 days post-transduction (Fig. 6E), thus indicating replicative arrest. Since *BRCA1* mutation carriers have one functional copy of *BRCA1*, we determined whether heterozygous loss of *Brca1* is sufficient to cause replicative arrest and/or onset of senescence in MOFs. Acidic β-galactosidase or senescence-associated β-galactosidase (SA-β-Gal) activity is a defining feature of senescence, used to distinguish senescent cells from quiescent cells and those in transient cell cycle arrest. *Brca1*-Het MOFs stained positively for SA-β-Gal activity 16 days post-transduction, while *Brca1*-WT MOFs showed little to no activity (Fig. 6F). Additionally, with loss of *Brca1*, there was decreased gene expression of *Mki67* and *Top2a* (Fig. 6G), which encode for Ki-67 and DNA topoisomerase II-α, both of which are highly expressed in cycling cells but strongly down-regulated in non-cycling cells and have been used as markers of proliferation (*39–41*). The onset of cellular senescence is associated with drastic morphological changes and cytoskeletal rearrangement, particularly that of vimentin, an intermediate cytoskeletal filament, often resulting in enlarged and flattened cells (*42*). Immunofluorescence staining of *Brca1*-KO MOFs for vimentin showed enlarged, flattened cell bodies compared to *Brca1*-WT MOFs (Fig. 6H).

Collectively, the upregulation of various CDK inhibitors, positive SA-β-gal staining, increased p16, decreased expression of the proliferation markers Ki-67 and Top2-α, and flattened, enlarged cellular morphology confirms the induction of cellular senescence with the loss of *Brca1* in MOFs.

### Secretory phenotype of senescent *Brca1* KO MOFs is enriched in components of senescence associated- P21 activated secretory phenotype

Cellular senescence is defined as a state of stable exit from the cell cycle that is accompanied by characteristic phenotypic changes to the cells, including the production of a bio-active secretome, referred to as the senescence-associated secretory phenotype (SASP). SASP is a dynamic, heterogeneous phenomenon that varies in composition, with time, and is biased by cell type and mode of senescence onset (*43*). Given the evidence that *Brca1*-KO MOFs have become senescent, we sought to identify the SASP signature of these cells. From the RNA-sequencing data, we identified the SASP signature of *Brca1*-KO cells to be enriched in cell adhesion molecules (*Icam1*, *Igfbp3* and *Igfbp7)* and matrix remodelling enzymes, such as *Mmp12* and *Mmp13* (Fig. 7A). The composition of SASP is governed by the activity of Trp53, either through positive regulation of NFкB, to produce a traditional strongly inflammatory SASP (NFкB SASP), or negatively regulate it to produce a unique non-inflammatory SASP enriched with modulators of the ECM (Trp53 SASP), in a mutually exclusively manner (*44*). RNA-sequencing data was analyzed using pre-established gene sets for Trp53 SASP and NFкB SASP (*44*) to determine the pathway governing the induction of SASP with the loss of *Brca1*. The gene module score of Trp53 SASP genes was significantly upregulated by *Brca1*-KO MOFs, while no significant differences were noted for those associated with NFкB SASP (Fig. 7B). Moreover, we observed significant increases in the expression of components and modulators of the ECM (*Col8a1*, *Efemp1* and *Lgals3*), proteinases and protease inhibitors (*Ctsk*, *Serpinb1a*, *Serpinb9b*, and *Serpinb6c*), and immunomodulatory genes (*B2m* and *C1qtnf3*), which have been identified as components of senescence associated p21-activated secretory phenotype (SA-PASP), which acts independently of Trp53 or NFкB (*45*) (Fig. 7C). Our data suggests that SASP of *Brca1*-KO MOFs is likely governed by the activities of Trp53 and p21.

**Fig 7:**
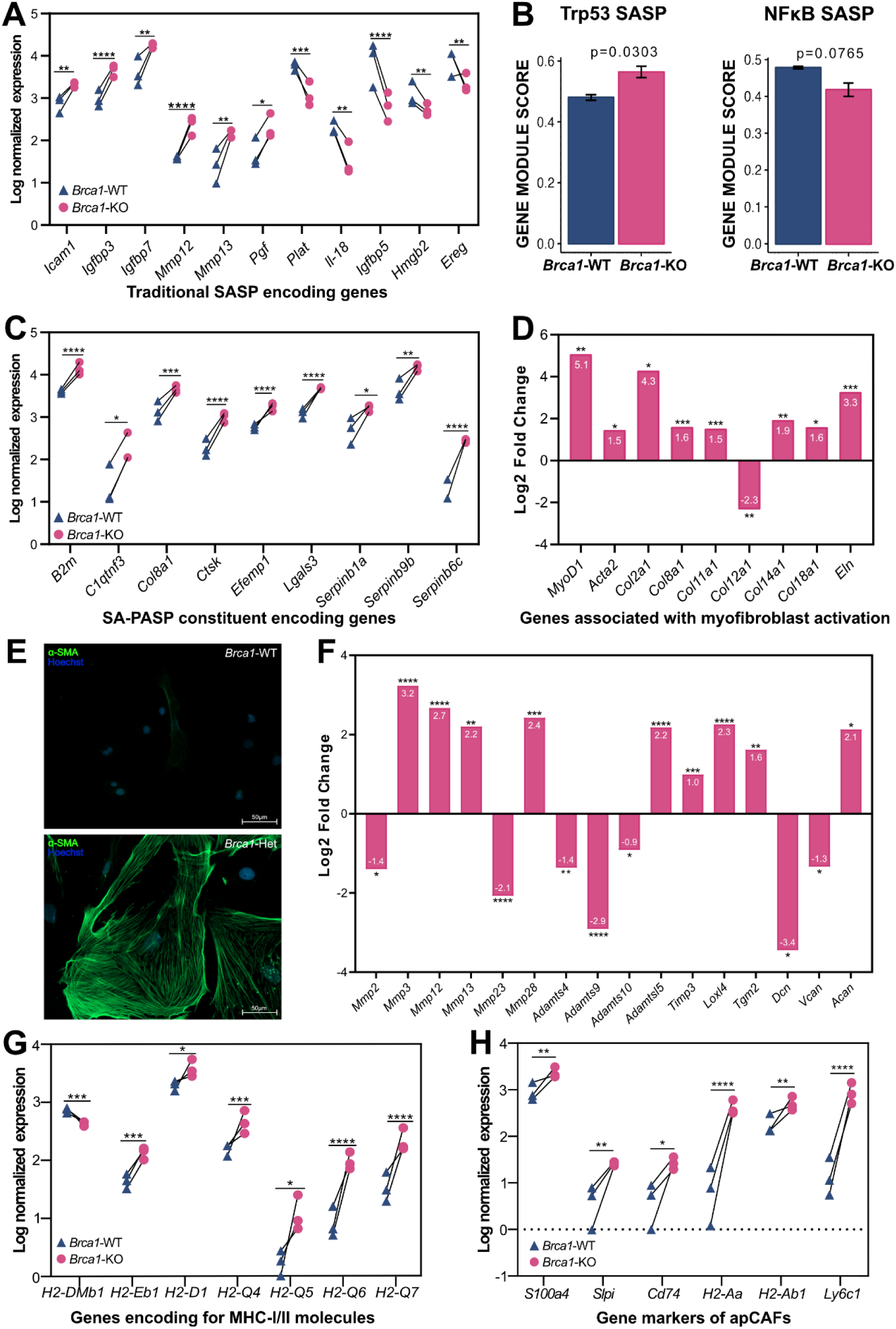
Loss of *Brca1* in MOFs induces a unique transcriptomic shift that results in a myofibroblastic apCAF-like state. *Brca1*^LoxP/LoxP^ MOFs were transduced with either AdCre (*Brca1*-KO) or AdEmpty (*Brca1*-WT) and were harvested 14 days later to extract RNA for RNA-sequencing or fixed for immunofluorescence staining of α-SMA (n=3). (**A**) SASP signature of *Brca1*-KO MOFs. (**B**) Overall SASP expression prolife of *Brca1*-KO MOFs is aligned with that driven by p53 but not NFκ-B. (**C**) Secretory phenotype of *Brca1*-KO MOFs is enriched with structural and enzymatic components of the ECM associated with SA-PASP. (**D**) *Brca1*-KO MOFs upregulate *MyoD1* and *Acta2*, markers associated with myofibroblast activation, that is characterized by upregulation of various collagen-encoding genes. (**E**) Immunofluorescence was used to visualize α-SMA stress fibers (green) in *Brca1*-Het MOFs, that were absent in *Brca1*-WT MOFs. Hoechst was used to visualize nuclei. Scale bars = 50 µm. (**F**) Loss of *Brca1* in MOFs results in upregulation of genes encoding for various enzymatic and non-enzymatic secreted components of the ECM that are associated with increased stiffness and dysregulation of the ECM. (**G**) Genes encoding for MHC-I and -II molecules were upregulated with loss of *Brca1*. (**H**) *Brca1*-KO MOFs upregulate gene markers of apCAFs. Differential gene expression is presented either as log normalized expression between *Brca1*-KO and *Brca1*-WT MOFs (**A**, **C**, **G**, **H**) or log2FC in *Brca1*-KO MOFs, compared to *Brca1*-WT MOFs (**D**, **F**), shown in white for each gene. Unpaired student’s t-test was performed using R. *= p < 0.05, **= p < 0.01, ***= p < 0.001, ****= p < 0.0001.

### *Brca1* KO in MOFs induces a myofibroblastic phenotype, indicative of ECM remodelling potential

Assessment of the secretome gene signatures of *Brca1*-KO MOFs revealed significant increases in the gene expression of numerous structural components of the ECM, notably *Col2a1, Col4a1*, *Col8a1*, *Col11a1*, *Col14a1*, *Col18a1*, and *Eln*. These were accompanied by upregulation of *Myod1*, encoding for MyoD, a transcription factor critical for the differentiation of myofibroblasts, as well as *Acta2*, which encodes for α-smooth muscle actin (α-SMA), a well-established marker of myofibroblasts (*46*, *47*) (Fig. 7D). Increased expression of these genes suggests that loss of *Brca1* in MOFs induces myofibroblast activation. Myofibroblasts are characterized by the formation of α-SMA stress fibers that are key to their contractile ability (*48*) Immunofluorescence staining revealed the formation of α-SMA stress fibers in both *Brca1*-KO MOFs (not shown) and *Brca1*-Het MOFs, but not *Brca1*-WT MOFs (Fig. 7E), thus confirming myofibroblast activation in these cells.

In addition to myofibroblast activation, there were marked changes in the gene expression of various ECM remodelling (*Mmp2*, *Mmp3*, *Mmp12*, *Mmp13*, *Mmp28*, *Adamts4*, *Adamts9*, *Adamts10*, *Adamtsl5*, *Serpinb1b*, *Serpinb1c*, *Serpinb9f*, and *Timp3*) and cross-linking enzymes (*Tgm2*), glycosaminoglycans (*Has2*, *Has3*), and proteoglycans (*Acan*, *Vcan*) that have been linked to increased stiffness and fibrotic remodelling of the matrix (Fig. 7F). These changes suggest that the loss of *Brca1* in ovarian fibroblasts may lead to dysregulation in the composition and structure of the surrounding ECM, mirroring the increased architectural changes in human BRCA1 Pre ovaries we identified (Fig. 3F).

### The loss of *Brca1* induces a transcriptomic shift that resembles the molecular profile of antigen-presenting cancer-associated fibroblasts

Since GOBP from GSEA identified upregulation of processes that mediate immune signalling and regulation, we assessed further the expression of immune modulatory molecules. Importantly, the gene expression of several components of the major histocompatibility complex (MHC) class-I (*H2-D1*, *H2-Q4*, *H2-Q5*, *H2-Q6*, *H2-Q7*) and -II (*H2-Aa*, *H2-Ab1*, *Cd74*, *H2-DMb1*, *H2-Eb1*) was significantly upregulated in *Brca1*-KO compared to *Brca1*-WT MOFs (Fig. 7G). While all cells express MHC-I, expression of MHC-II, previously thought to be restricted to “professional” antigen-presenting cells (*49*), can also be expressed by a novel subtype of cancer-associated fibroblasts (CAFs), referred to as antigen-presenting CAFs (apCAFs). Interestingly, we observed significant upregulation of several gene markers of apCAFs (*50–53*), namely, *S100a4* (log2FC=1.31), *Slpi* (log2FC= 2.49), *Cd74* (log2FC=2.49), *H2-Aa* (Log2FC= 5.37), *H2-Ab1* (Log2FC=1.41) and *Ly6c1* (logFC=5.66) in *Brca1*-KO MOFs (Fig. 7H), indicating a potential shift towards an apCAF-like state, at least at the transcriptomic level.

## Discussion

While the cancer risk in *BRCA+* carriers has been extensively studied in the context of epithelial cell transformation (*4*, *5*), there is much less known about the impact of BRCA dysfunction in non-epithelial cells. Stromal hyperplasia in *BRCA+* ovaries (*54*, *55*) and hyperactivity in skin fibroblasts containing a *BRCA1* epimutation (*29*) have been reported, with BRCA haploinsufficiency resulting in replication stress due to defects in stalled replication fork repair (*29*, *56*). This study provides insight into other mechanisms by which *BRCA1*/2 haploinsufficiency may have pathogenic outcomes. Given the links between *BRCA* mutation, early loss of ovarian function and increased cancer risk, we investigated whether *BRCA* mutation contributes to the development of ovarian fibrosis. While we and others have shown that ovarian fibrosis occurs in an age-dependent manner in both humans and mice (*13*, *24*, *57*), this study is the first to uncover that ovarian fibrosis is present in the ovaries of cancer-free *BRCA+* carriers at an earlier age than in non-carriers. Exploring the possible mechanisms leading to this fibrosis using *Brca1*-mutant MOFs revealed increased DNA damage associated with myofibroblast activation, senescence and phenotypic changes resembling CAFs.

While the definition of fibrosis can vary depending on the tissue type, we have noted here and in a previous study (*13*) that ovarian fibrosis in humans is characterized by the orientation of collagen fibers rather than the accumulation of collagen. Collagen composition (amount and type), crosslinking, density (fibril packing and size) and orientation (alignment and crimp) all encompass the global collagen architecture within a tissue (*58*). Our PSR-POL analysis identified increased alignment of collagen fibers within the ovarian cortex, however, this alongside collagen content only accounted for a small proportion the collagen architecture and its contribution to fibrosis. We therefore expanded our knowledge of the collagen landscape within the ovarian cortex by performing single-fiber analysis using the FibroNest digital pathology platform. This analysis confirmed the results of the PSR-POL experiment with *BRCA*+ carriers exhibiting higher fibrosis scores than non-carriers although there was variability between samples. The outcomes are also in agreement with our previous finding that postmenopausal ovaries had significantly higher degrees of fibrosis than premenopausal ovaries (*13*).

*BRCA+* carriers exhibited a range of fibrosis scores and had a distinct fibrosis phenotype. Two traits that were exclusive to *BRCA+* carriers were lowered tortuosity and orientation. Lower tortuosity, which indicates fewer curves and twists within the collagen fibers, has been shown to reduce the strength and elasticity of a tissue, both of which are apparent in fibrosis (*58*). Collagen orientation is also lower by FibroNest analysis, however, orientation in this analysis refers to the fiber alignment in relation to the north axis of the image while PSR-POL dominant direction analysis identifies collagen alignment in relation to the dominant direction. Therefore, since these two analyses of orientation are measuring different types of orientation, we can conclude that age and BRCA mutation alter the orientation and alignment of collagen fibers. This collagen linearity is a contributor to collagen architecture, and therefore it is logical that the global collagen architecture score was significantly different from the age-matched controls. It should be noted that for most of the scores, the WT Pre ovaries had low fibrosis scores, the WT Post ovaries had high scores and the BRCA Pre ovaries were intermediate. This pattern may indicate a different mechanism of fibrosis development in young *BRCA+* carriers. Whether this altered structure creates a niche that is more supportive of ovarian cancer growth remains to be determined. Given the high degree of variability between *BRCA*+ patients, we speculate that the higher values reflect more fibrotic ovaries that are forming a tumor permissive niche. The penetrance, type, and resulting function of *BRCA* mutations are also variable and could account for this high degree of variability within patient samples (*59*). While all *BRCA* mutations in this cohort were pathogenic, not all *BRCA*+ carriers resembled the WT Post ovaries or even other ovaries within the BRCA Pre group, which we speculate might reflect differences in risk seen between pathogenic variants.

One limitation of the collagen profiling in fixed human tissue is that it reflects a single point in time, and few conclusions can be drawn regarding the mechanism of fibrosis development. Therefore, we used a *Brca1* inducible knockout model to study the effects of heterozygous and homozygous deletion of *Brca1* in MOFs. The deletion of *Brca1* from primary cultures of MOFs led to the accumulation of dsDNA damage and induced the onset of cellular senescence characterized by an enlarged, flat morphology, increased expression of cell cycle inhibitors, specifically *Cdkn2a*, positive staining for SA-β-gal activity, and decreased expression of markers of proliferation. These alterations were accompanied by the induction of a unique myofibroblastic, antigen-presenting CAF-like phenotype that was characterized by increased expression of *Acta2*, formation of α-smooth muscle actin stress fibers and upregulation of a multitude of structural and enzymatic components of the ECM, as well as those encoding MHC-II molecules and other markers of apCAFs. These results suggest that loss of BRCA1 function in ovarian fibroblasts has the potential to create a pro-fibrotic niche, a microenvironment that has been widely implicated in supporting tumor growth in other tissues (*60*, *61*). We hypothesize that an earlier onset of fibrosis in the ovarian stroma may contribute to premature ovarian aging, and the early onset of ovarian cancer observed in *BRCA+* carriers (*19*, *21*).

Given that BRCA1 is best known for its integral role in preserving genomic integrity, it is not surprising that heterozygous loss of *Brca1* results in the accumulation of dsDNA damage, and that the transcriptomic profile of *Brca1*-KO MOFs is indicative of impaired dsDNA repair with reduced expression of DNA repair genes. It has been proposed that the accumulation of DNA damage due to the loss of BRCA1 function in epithelial cells likely evokes a cellular stress response through the activation of Trp53, initially resulting in cell cycle arrest, followed either by apoptotic clearance or induction of cellular senescence to prevent propagation of genetically compromised cells (*38*, *62*). The cellular response of MOFs to the loss of *Brca1*, characterized by initial cell death through apoptosis followed by the onset of senescence in surviving cells, is consistent with the proposed activation of Trp53 in response to irreparable DNA damage.

The onset of cellular senescence, a key hallmark of aging, is defined as a stable state of cell cycle arrest that is coupled with characteristic phenotypic and metabolic changes triggered by stressors compromising genomic stability (*43*). The development of senescence is initially dominated by p53 activity that induces transient cell cycle arrest. This is followed by activation of the p16/Rb pathway, which is characterized by irreversible cell cycle arrest, elevated expression of p21 and p16, SA-β-gal activity, decreased expression of Ki67 (*Mki67*) and production of SASP (*63*), all of which are exhibited by *Brca1*-KO MOFs, thus further validating that loss of BRCA1 function in ovarian fibroblasts results in the induction of senescence.

The production of SASP is a key pathogenic feature of senescent cells due to its ability to permanently alter the tissue microenvironment. Though the composition of SASP is highly variable and diverse, it is often enriched with pro-inflammatory cytokines, chemokines and interleukins (*44*). Surprisingly, the secretome reflected by the transcriptome of *Brca1* KO MOFs was entirely devoid of the traditionally inflammatory components, but was heavily enriched with various ECM components, including structural and matricellular proteins, remodelling and cross-linking enzymes, glycosaminoglycans, and proteoglycans. This suggests that the SASP production of *Brca1*-KO MOFs is likely independent of NFкB and may be governed by the activities of Trp53 and p21, which contribute to an alternative mechanism that induces senescence and is more associated with ECM remodelling.

Interestingly, the induction of this unique non-inflammatory secretory phenotype enriched with ECM modulators substantially overlaps with the observed myofibroblastic phenotype of *Brca1*-KO MOFs. Induction of a myofibroblastic phenotype in *Brca1*-KO MOFs is also consistent with the phenotype reported for dermal and stromal breast fibroblasts heterozygous for *BRCA1* (*29*, *64*). Myofibroblasts, which are contractile cells typically associated with upregulation of fibrillar collagens I and III that form the structural framework of the ECM, are indispensable for wound healing and maintaining tissue homeostasis. While we did not observe upregulation of genes encoding for either collagen I or III, the expression of several others was increased in *Brca1*-KO MOFs, including those encoding for collagen II, XI and XIV. Collagen II expression is native mostly to cartilaginous tissue, where microfibrils of collagens II and XI associate to form dense and rigid supramolecular collagen fibrils (*65*). Collagen XIV, a fibril-associated collagen that provides increased stability to the ECM is present in areas of mechanical stress, such as in bleomycin-induced fibrotic lesions in the lungs (*66*). The shift to a transcriptome reflecting enhanced fibrosis-generating capacity is also indicated by dysregulated expression of proteoglycans, such as reduced expression of decorin (encoded by *Dcn*) that functions as a structural spacer by simultaneously interacting with various collagens to limit the anisotropic assembly of collagen fibrils, a defining characteristic of fibrosis (*67*).

Other proteoglycans, such as aggrecan and versican, which interact with and stabilize collagen fibrils to create a network of collagen. According to the human protein atlas, while versican is native to the ovarian ECM, aggrecan is not, it is almost exclusively expressed in cartilaginous tissues, where it interacts with collagen II to generate a uniquely stiff and dense network that is resistant to degradation (*68*). We observed upregulation of *Acan*, encoding for cartilage-specific proteoglycan aggrecan and downregulation of *Vcan*, encoding for ovarian native proteoglycan versican. Interestingly, we noted decreased expression of aggrecanase encoding genes, *Adamts9* and *Adamts4*, which may facilitate the increased deposition of aggrecan-collagen II aggregates. Covalent cross-linking of collagen fibrils limits the flexibility of the collagen-proteoglycan network (*69*). As such, excessive cross-linking is a major driving force of age- and fibrosis-associated tissue stiffness and mechanical stress (*70*, *71*). This is supported by increased expression of cross-linking enzymes, such as transglutaminase 2 and members of LOX and LOXL families, in cases of idiopathic pulmonary fibrosis (*72*, *73*). Consistent with this, *Brca1*-KO MOFs upregulated genes encoding for cross-linking enzymes Loxl4 and transglutaminase 2 (encoded by *Loxl4* and *Tgm2*, respectively), that are responsible for crosslinking fibrils of collagens II/II, II/XI, XI/XI and collagen-elastin fibers (*74–76*). Collectively, the transcriptome predicts a secretory phenotype of *Brca1*-KO MOFs that is clearly characterized by dysregulated expression of various ECM components, including structural proteins, cross-linking and remodelling enzymes, as well as glycoproteins. This conveys the strong remodelling potential of myofibroblastic *Brca1*-KO MOFs that may prime dysregulation in the composition, structure and stiffness of the ovarian ECM.

Elimination of the myofibroblastic phenotype is just as important as the activation of myoblasts in maintaining homeostasis. The transcription factor MyoD is a critical switch for the differentiation and de-differentiation of myofibroblasts. Sustained upregulation of MyoD precedes that of *Acta2* during myofibroblast differentiation, and declining levels of MyoD precede those of α-SMA as myofibroblasts de-differentiate into fibroblasts (*77*, *78*). Myofibroblasts with sustained, elevated MyoD fail to undergo de-differentiation and display resistance to apoptosis, whereas genetic silencing of MyoD restores susceptibility to apoptosis (*78*). The accumulation of myofibroblasts, either due to impaired de-differentiation or resistance to apoptosis, has been implicated in the pathology of fibrotic disorders (*77*). As such, elevated expression of *Myod1* in *Brca1*-KO MOFs may be a mechanism driving the persistent myofibroblastic phenotype.

The expression of the non-classical MHC-1b molecule, H2-q, has been shown to confer senescent cells with the ability to evade immune clearance by inhibiting the activity of NK and CD8+ T-cells by rendering them functionally exhausted through the inhibitory receptor NKG2a (*79*). As such, upregulation of H2-q6/7 in *Brca1*-KO MOFs may confer the ability to evade clearance by both innate and adaptive immune cells. Additionally, the expression of MHC-II encoding genes (*H2-Aa, H2-Ab1, H2-Eb, and Cd74*), that is typically restricted to “professional” antigen-presenting cells, is indicative of antigen presenting capacity of *Brca1* KO MOFs, which may allow these senescent myofibroblasts to modulate adaptive immune activity.

Growing evidence attributes physiological aging and age-associated tissue fibrosis to the accumulation of senescent myofibroblasts (*80*, *81*). Ovarian aging is accompanied by substantial age-associated changes to the ovarian stroma, specifically, increased stiffness due to the development of fibrosis (*82*, *83*). These age-associated changes may be attributed to the presence of α-SMA+ and p16+ cells in the stroma, that likely correspond to myofibroblastic and senescent cells, respectively (*84*, *85*). In *Brca1*-KO MOFs, the combined onset of senescence and myofibroblast differentiation, both of which can confer resistance to apoptosis, suggests the potential to facilitate the persistence and accumulation of these senescent myofibroblasts in the ovary. Moreover, the expression of MHC-II and non-classical MHC-I genes by non-immune cells, such as *Brca1*-KO MOFs, is considered a gene signature for aging that has been observed in SASP-producing fibroblasts unique to aged fibrotic ovaries (*57*). Given the parallels observed between the transcriptomic profile of *Brca1*-KO MOFs and age-associated features of the ovarian stroma, it appears that the loss of *Brca1* reprograms the transcriptome of ovarian fibroblasts towards a state reminiscent of aging. We propose that the accumulation of these senescent, antigen-presenting myofibroblasts in the ovary is a possible mechanism for the creation of an ovarian niche supportive of tumor growth in *BRCA+* carriers. Additionally, diminished replicative capacity of ovarian stromal fibroblasts and increased stiffness of the ovarian ECM, which are likely to impede various aspects of ovarian function (*83*, *86*), provide an explanation for the premature ovarian aging observed in *BRCA1* mutation carriers.

## Materials and Methods

### Patient Sample Collection

A cohort of human ovarian samples was accrued from collaborators at the Centre de recherche du Centre Hospitalier de l’Université de Montréal (CRCHUM, Montreal, Quebec, Canada) and the OCRC Tumour BioTrust Collection (RRID:SCR_022387; Philadelphia, Pennsylvania, USA) and the Ottawa Ovarian Cancer Biobank. The study was approved by all three institutional ethics committees: the Comité d’éthique de la recherche du CHUM (CÉR-CHUM) #2005-1893: BD 04.002, IRB Protocol Number 702679 for the BioTrust Collection, and the Ottawa Health Science Network Research Ethics Board protocol 20180168-01H. Ovaries were collected following oophorectomies, fixed in formalin and paraffin embedded (FFPE). Ovaries of patients without *BRCA* mutations were removed for uterine and/or cervical pathologies. All ovaries were assessed by pathologists and considered structurally normal and non-cancerous. Tissues were stored in respective biobanks until retrieval at which time they were sectioned (5 μm) and shipped. This cohort included 53 samples from both pre- and post-menopausal *BRCA1 or BRCA2* mutation carriers and non-carriers. The patients range in age from 36-83. Patient information can be found in Supplementary Table 1.

### PSR Staining and Imaging

Histopathologic assessment of human ovaries was performed on 5 μm sections of FFPE tissue. All sections were stained with picrosirius red (PSR) at the Louise Pelletier Histology Core Facility at the University of Ottawa. Slides were imaged at 40X under polarized light (PSR-POL) using a Zeiss AxioObserver 7 Inverted Microscope (with tunable polarizer and analyzer; Zeiss; CBIA Core Facility at the University of Ottawa). Regions were selected across the ovarian cortex by one investigator and then these regions were located and imaged by another investigator to reduce bias. A four-tile area was taken for each image and five regions imaged across the ovarian cortex for each sample to get a representative range of collagen deposition. Collagen quantification was performed on polarized images using Fiji (ImageJ). For each image, a region of interest (ROI) of 100 mm × 100 mm was created and positioned 50 mm below the surface of the ovary to ensure the ovarian surface epithelium was excluded from the analysis.

PSR-POL analysis was performed as described previously (*57*). Briefly, a HUE program was run to determine the collagen abundance as it filters the images for number of pixels that are at wavelengths related to the 4 colours seen in PSR-POL (green, yellow, orange and red). Definition of hue was described previously (*87*) by wavelength: green, 52 to 128 nm; yellow, 39 to 51 nm; orange, 10 to 38 nm; red, 2 to 9 and 230 to 256 nm. The five regions across the ovarian cortex were run through the HUE program individually, and the number of pixels for each fiber colour was calculated as a percentage of the total ROI area in pixels. The geomean was calculated across the five regions per sample and a t-test was run between the pre- and postmenopausal groups. Total collagen fiber content includes all hue determined pixels.

PSR-POL images were also used to quantify the alignment of collagen fibers within the ovarian cortex. The OrientationJ package was used to run the Dominant Direction program for collagen coherency. The directionality of collagen fibers in a selected ROI was assessed to determine whether a directionality was predominant and what percentage of fibers were running in said dominant direction (*88*). With this program, the resulting coherency coefficient can range from 0 (no preferential orientation of fibers) to 1, the most coherent orientation of the fibers. The five regions across the ovarian cortex for each sample were run through the program individually, the average was calculated, and an unpaired t-test was performed to compare the pre- and postmenopausal groups.

### Histological Quantification of Ovarian Fibrosis by Digital Pathology

Collagen was also profiled with digital pathology imaging to quantify the broad features of the fibrotic phenotype in human ovarian cortex. Brightfield images of FFPE ovarian cortex stained with PSR were taken at 40X using the AxioScan 7 (Zeiss) and uploaded to the FibroNest quantitative digital pathology platform (PharmaNest, Princeton, NJ). ROI were selected within the ovarian cortex beginning 30 μm below the surface and extending 300 μm in depth (see Supplementary Fig. 3 for raw images). The FibroNest platform quantified three levels of fibrosis: collagen amount and structure (12 traits), collagen fiber morphometry (13 traits) and collagen architecture (7 traits), which all contributed to the overall histological phenotype of fibrosis. The analysis was further subdivided to quantification of fine and assembled collagen fibers. Fine collagen fibers are defined as having <295 branches, are less complex and usually indicate the beginnings of fibrosis. Assembled fibers are highly complex with >295 branches and are indicative of established fibrosis. Each of the aforementioned traits were used to create a histogram where the trait is described by up to 7 quantitative fibrosis traits (qFTs) to complete statistical quantification. Upwards of 300 qFTs are assessed; only those that describe most of the variability between groups (principal qFTs, 177 in total) were used to create the composite fibrosis scores. Significance of heatmap scores between analysis groups was determined using Welch’s t test and p < 0.05. Significance of fibrosis scores were determined using a one-way ANOVA with Tukey’s post-hoc multiple comparisons test.

### Experimental Animals

*Brca1*^LoxP/LoxP^ [FVB;129-*Brca1*^tm2Brm^EYFP] conditional knockout mice, bearing LoxP sites in introns 4 and 13 of the *Brca1* gene, were generated by breeding *Brca1*^tm1Brn^ mice to B6.129X1-Gt(ROSA)26Sor^tm1(EYFP)Cos^/J mice. The resulting mice bear a LoxP-flanked STOP sequence followed by the Enhanced Yellow Fluorescent Protein (EYFP) gene and its associated polyA tail. *Brca1*^LoxP/LoxP^ were crossed with FVB/N mice to generate *Brca1*^LoxP/+^ mice. All experiments were performed according to the Guidelines for the Care and Use of Animals established by the Canadian Council on Animal Care using a protocol (OHRIe-3522) approved by the University of Ottawa Animal Care Committee, and with adherence to the ARRIVE guidelines.

### Primary murine ovarian fibroblast (MOF) isolation and culture

*Brca1*^LoxP/LoxP^ or *Brca1*^LoxP/+^ mice between the ages of 6-10 weeks were euthanized via CO_2_ asphyxiation and dissected to collect both ovaries. Seven to ten mice were used for each isolation. Freshly collected ovaries were stored in cold phosphate-buffered saline (PBS; Gibco; #14190-144) on ice during collection. Using a stereomicroscope (Leica WILD M3C), any remaining adipose tissue as well as the ovarian bursa were removed. Ovaries were then washed with cold sterile PBS and punctured repeatedly with 26- and 27-gauge needles to release the oocytes and granulosa cells from growing follicles. The ovaries were then gently patted flat using a blunt probe to ensure the elimination of as many granulosa cells as possible. Residual ovarian tissue was rinsed with fresh PBS several times and then transferred to enzymatic cocktail I [8mg/mL collagenase III (StemCell technologies; #07423) and 1500 kunitz units DNase I (Qiagen; #79256) in Hank’s balanced salt solution (HBSS, Gibco;14170-112); 1mL enzymatic cocktail I for 2 ovaries]. The ovaries were minced using sterile dissection scissors to aid the digestion of the ECM proteins and incubated in enzymatic cocktail I at 37°C for 15 mins on a rotary shaker, gently mixing every 5 mins. The dissociated tissue was centrifuged at 0.4g for 5 mins and resuspended in enzymatic cocktail II [1500 kunitz units DNase I in 0.25% trypsin (Gibco; #25200-056); 1mL enzymatic cocktail II for 2 ovaries] to further break down the connective tissue proteins. Following a 10-minute incubation at 37°C on a rotary shaker, trypsin was inactivated by the addition of warm MOF media [filtered 1xDMEM (Corning; #10-013-CV) containing 10% FBS (Gibco; #12483-020 and Corning; #35-077-CV), 1% PenStrep (Gibco; # 15140-122) and 4 ng/mL of FGF (R&D system; #3139-FB)]. The tissue mixture was passed through a pre-wetted 40 µm nylon mesh cell strainer (ThermoFisher Scientific; #22363547) to remove any oocytes and tissue debris. The filtered cell suspension was then centrifuged at 0.4g for 5 mins and resuspended in 1mL of warm MOF media. The cell suspension was immediately plated in a pre-warmed fibronectin-coated T25 flask [Tissue culture plates and coverslips were coated with 20µg/mL human plasma fibronectin (Sigma-Aldrich; #F0895-1MG) in HBSS (Gibco, #14170-112) at a concentration of 2µg/cm^2^ and kept at room temperature (RT) for an hour, prior to being rinsed with PBS and drying for 2-3h in a tissue culture hood] and cultured at 37°C in hypoxic conditions (5% O_2_, 5% CO_2_). MOFs were passaged 7-10 days later, upon reaching confluence.

### Transduction of MOFs with adenovirus

Primary *Brca1*^LoxP/LoxP^ and *Brca1*^LoxP/+^ MOFs were transduced with either recombinant adenovirus expressing cre recombinase, Ad5CMVCre (AdCre; Vector Development Laboratory) or recombinant empty adenovirus, Ad5CMVEmpty (AdEmpty; Vector Development Laboratory) or no virus (control) at passage 1 (p1). MOFs at confluence (p0) were washed with PBS, detached using 0.05% trypsin (Gibco; #25300-054), centrifuged at 0.4g for 5 mins at RT, resuspended in 2ml of warm serum-free MOF media (filtered 1xDMEM containing 1% PenStrep) and counted using Cell Countess^®^ II (Life technologies; #AMQAX1000). Double the number of MOFs were transduced with AdCre than with AdEmpty to account for substantially reduced cell count following *Brca1* KO-induced cytotoxicity. Cells were incubated in poly-L-lysine (1µg/mL; Sigma Aldrich; #P9155) for 15 mins at RT prior to exposure to adenovirus to increase the efficiency of virus transduction. MOFs were transduced with either AdCre or AdEmpty in suspension at 50 multiplicity of infection (MOI) for 3.5-4 hours at 37°C in hypoxic conditions. Transduced and control cells were immediately plated on fibronectin-coated 6-well plates. After 4 hours, once the cells have attached, adenovirus-containing media was removed and replaced with warm MOF media. Cells were then allowed to recover from transduction for up to 16 days at 37°C in 5% O_2_ and CO_2_. The resulting primary cultures of transduced cells were defined by their BRCA1 status: *Brca1*^LoxP/LoxP^ cells transduced with AdCre will be hereafter referred to as *Brca1*-KO; *Brca1*^+/-^ will hereafter be referred to as *Brca1*-Het. *Brca1*^LoxP/LoxP^ MOFs transduced with AdEmpty will be referred to as *Brca1*-WT.

### EYFP monitoring

EYFP expression by the Rosa26 locus was used to track *Brca1* deletion and to ensure that the cells prior to RNA/DNA/protein extraction were *Brca1* negative cells (i.e., EYFP positive cells). EYFP expression was monitored in *Brca1*-KO and *Brca1*-Het MOFs for 14 days, starting 48h after transduction. Fluorescence was detected using excitation and emission spectra of 500nm and 539nm, respectively, using a Biotek cytation 5 imaging reader. The transducion efficiency varied between 70-80%.

### Apoptosis assay

RealTime-Glo Apoptosis Assays (Promega; JA1000) were used to measure apoptosis in control, *Brca1*-WT, and *Brca1*-KO MOFs over a span of 14 days, starting 4 hours after exposure to virus. MOFs (10,000 cells for control and AdEmpty and 12,000 cells for AdCre) were plated and transduced in flat, clear-bottom 96 well ImageLock plates (Sartorius, #4379). Apoptosis assay kit components were added to the cell culture media at 1x concentration, as per the manufacturer’s protocol, at the end of virus infection. Luminescence produced by functional luciferase was measured every 24 hours using a Biotek cytation 5 imaging reader at a gain of 100, 150 and 200. Raw data (gain 100) was normalized using mean of negative control (no cells with kit components). Two-way ANOVA was applied using GraphPad Prism.

### RNA and genomic DNA (gDNA) extraction

RNA, gDNA and protein were extracted using Allprep DNA/RNA/protein mini kit (Qiagen; # 80004) as per the manufacturer’s protocol. Extracted RNA was used for RNA-sequencing and to synthesize cDNA using iScript Reverse Transcription Supermix, as per manufacturer’s protocol (Bio-Rad; #1708841). A Bio-Rad T100 thermal cycler was used to perform reverse transcription, and cDNA was then used for quantitative polymerase chain reaction (PCR). Extracted gDNA was used to confirm cre-mediated recombination of floxed *Brca1*.

### Polymerase chain reaction and gel electrophoresis

Each PCR to detect recombination of *Brca1* was prepared using milli-Q H_2_O to a final volume of 10 µL, with 1x concentration of Extract-N-Amp (stock at 2x; Sigma XNAT2R), 0.40-0.5 µM of each primer (stock at 10 µM), and 2 µL gDNA. Primers (*Brca1*-intron4_F & -intron13_R, TATCACCACTGAATCTCTACCG and TCCATAGCATCTCCTTCTAAAC, respectively, yield a 600bp amplicon; *Brca1*_exon11_F&R, ATCAGTAGTAGAAATCCAAGCCCACC and TGCCTCTCCCAGCATTGTTAG, respectively, yield a 592bp amplicon). Primers were obtained from Invitrogen. PCR amplification was carried out in a Bio-Rad T100 thermal cycler as follows: initial denaturation at 94°C for 3 mins followed by denaturation, annealing and extension at 94°C, 56°C, and 72°C, respectively, for 30 seconds each. This sequence was repeated 30 times, which was followed by a final extension step at 72°C for 10 mins. PCR products were stored at 4°C and separated on 1.5% agarose gels (1% SafeRed) in Tris acetate-EDTA buffer for 45 mins at 110 volts.

### Quantitative polymerase chain reaction (qPCR)

*Brca1*F&R primers (GAAGCGGGGAGAATGGACTC and CTCTCAAATCGTCCCTCTGTCA, respectively; 139bp amplicon; annealing temperature: 59°C) used for qPCR were designed and validated using pooled cDNA from MOFs. Samples for qPCR were prepared using SsoAdvanced Universal SYBR^®^ green (Bio-Rad, #1725274), as per the manufacturer’s protocol. *Ppia*, *Gapdh*, *Hrpt1* and *Act-β* were used as housekeeping genes to normalize raw data collected using an ABI 5700 Fast Real-Time PCR system. Data was analyzed using the delta-delta-CT (ΔΔCT) method to compare expression of *Brca1* in *Brca1*-KO MOFs relative to *Brca1*-WT MOFs. Student’s t-test was performed, using GraphPad Prism 9. Difference in gene expression is presented as relative (log2) fold change.

### Western Blot

Protein was collected by cell lysis with RIPA buffer, 6 μM NaVO3, 10 ug/mL pepstatin and 10 μM PMSF. The proteins were denatured at 100°C for 5 min and loaded into a 15% tris-glycine gel. The gel ran for 60 min at 100 mA in 1X running buffer. Protein was transferred for 90 min at 100V into a PVDF membrane (ThermoFisher; P188520) with transfer buffer containing 20% methanol. The membrane was incubated in the primary antibody (p16; Abcam ab211542), diluted with 5% bovine serum albumin (BSA, BioShop; ALB001) at 1:1000 overnight at 4°C on a shaker. The membrane was washed, and secondary horseradish peroxidase antibody was diluted in BSA for 60 min at room temperature on a shaker. The membrane was washed and imaged with western enhanced chemiluminescence substrate (Bio-Rad; 1705061) in the BioRad ChemiDoc imaging system.

### Immunofluorescence

*Brca1*-WT, *Brca1*-KO and *Brca1*-Het MOFs were plated at a seeding density of 3 × 10^5^ for WT cells and 6 × 10^5^ for KO and Het cells/well (per 2mL media) on fibronectin-coated coverslips. Double the number of MOFs were transduced with AdCre to account for higher cell loss following *Brca1* knockout. Day 14 after transduction, conditioned media were removed from the MOFs and cells were washed with PBS. MOFs were fixed using 4% paraformaldehyde for 20 mins at RT, then permeabilized and blocked using 0.2% Triton x-100 (15 mins) and 10% goat serum (1h). Permeabilized and blocked cells were incubated with primary antibodies for either 1h at RT or overnight at 4°C. Primary antibodies were diluted in 10% goat serum at the following concentrations: rabbit anti-vimentin (Abcam; ab45939; used at 1 µg/mL), mouse anti-α smooth muscle actin (Invitrogen; #14-9760-82; used at 1 µg/mL), rabbit anti- γH2AX (Cell Signaling; #2577L; used at 1:100 dilution). Following three 5-minute washes with 1X PBS, cells were stained with either anti-mouse or anti-rabbit secondary antibodies (diluted in 10% goat serum) for 1h at RT, at the following concentrations: Alexa Fluor 647 donkey anti-mouse (ThermoFisher Scientific; #A-31571), at 1:1000; Alexa Fluor 488 donkey anti-mouse (ThermoFisher Scientific; #A-21202) at 0.2 µg/mL; Alexa Fluor 594 goat anti-rabbit (ThermoFisher Scientific; #A-11012) at 1:1000; Alexa Fluor 647 goat anti-rabbit (ThermoFisher Scientific; #A-21244), Alexa Fluor 488 goat anti-mouse (Invitrogen; #A10680) at 1:1000; Alexa Fluor 488 goat anti-rabbit (Invitrogen; #A11034) at 1:1000. Nuclei were then stained using Hoechst (1:250) in 1X PBS for 5 minutes, followed by three 5-minute washes with 1X PBS to remove excess secondary antibodies. Coverslips were mounted onto Superfrost plus microscope slides (ThermoFisher Scientific; #12-550-15), using either Epredia Immu-Mount (ThermoFisher Scientific; #9990402) or ProLong diamond antifade mountant with DAPI (ThermoFisher Scientific; #P36971). Images were acquired using an Axioskop2 MOT fluorescence microscope.

### Acidic senescence associated-β-galactosidase activity staining

*Brca1*-Het and *Brca1*-WT MOFs were stained using the senescence associated β-galactosidase activity assay (SA-β-Gal stain). Sixteen days post-transduction, MOFs were washed with PBS and fixed with 0.5% glutaraldehyde in PBS (ThermoFisher Scientific, #BP2548-1) for 15 mins at RT. Fixed MOFs were rinsed and stored at 4°C in PBS overnight. The following day, cells were washed with 1mM MgCl_2_ in PBS at pH5.5 for 10 mins, twice, at RT. Cells were then incubated in pre-heated (37 °C) X-gal staining solution comprising of 5% 20X KC solution (100mM potassium ferricyanide, 100mM potassium ferrocyanide trihydrate, in 1x PBS) and 2.5% of 40X X-Gal solution (1mg/mL x-gal from Bioshop, #XGA001 in N,N-Dimethylformamide) in 1mM MgCl_2_ (pH 5.5) at 37°C, in the dark for 35-45 minutes. The cells were then washed with 100mM Tris at pH 8.0 for 1 min at RT three times to stop the reaction. Cells were imaged immediately in 100mM Tris using an EVOS core AMEX-1200 transmitted-light inverted imaging system (Invitrogen; #12-563-649).

### RNA sequencing

*Brca1*-WT and *Brca1*-KO cells were generated in three independent replicate experiments. RNA was extracted 2 weeks after transduction (AllPrep DNA/RNA/Protein minikit, Qiagen) and quantified using a Nanodrop Spectrophotometer ND-100 (Nanodrop Technologies Inc.). RNA integrity was assessed using a Bioanalyzer (Agilent Technologies). Each sample containing at least 500ng of RNA was sequenced by Genome Québec on a NovaSeq6000 sequencer. RNA-seq data was aligned using Kallisto and differential gene expression analysis was performed using DESeq2 (R Project) (*89*). Gene sets were either acquired from MSigDb or adapted from literature or custom designed and scored using Singscore (R Project), which applies a scoring method proposed by Foroutan et al. (*31*) and Bhuva et al. (*32*) to generate a gene module score using rank-based statistics to analyze each sample’s gene expression profile and score the expression activities of gene sets at a single-sample level. As such, the gene module score for a given gene set is a rank-based average of all genes within that gene set. Gene sets acquired from MSigDb: “DNA double strand break repair”, “DNA double strand break response”, “Inhibition of replication initiation via Rb-E2f1”. Gene lists for “Senescence core genes” and “Traditional SASP encoding genes” was acquired from Landry et al. 2022 (*57*). Gene sets “Trp53 SASP” and “NAFкB SASP” were acquired from Lopes-Paciencia et al. 2019 (*44*). Datasets generated by Sturmlechner and colleagues 2021 (*45*) were used to custom build “Senescence-associated p21 activated secretory phenotype (SA-PASP)” gene list. Gene list “Markers of antigen presenting (ap)CAFs” was adapted from Elyada et al 2019, Friedman at al 2020, and Cords et al 2023 (*51–53*). Normal student’s t-tests were used for all statistical analyses (R Project). Genes with p-value > 0.05 were not considered in this study.

## Supporting information

Brca Manuscript_Supplemental Figures

## Acknowledgments

We are grateful to Julee Pauling and Sylvie Horne, our patient partners, for their valuable input into the goals of this project. We thank Dr. Nhung (Rose) Vuong for helpful discussions and Dr. Sarah Nersesian for critical review of the manuscript. We would also like to acknowledge the Cell Biology and Image Acquisition Core (RRID: SCR_021845) funded by the University of Ottawa and the Canada Foundation for Innovation, the Animal Care and Veterinary Services Core Facility at the University of Ottawa, and the Louise Pelletier Histology Core Facility at the University of Ottawa. We are grateful to PharmaNest for completing the FibroNest analysis.

## Funding

This work was supported by a grant from the Canadian Institutes of Health Research (E-419853) and a generous donation from the Carol Annibale Ovarian Cancer Foundation. We also acknowledge and honor the late Ms. Margaret Craig for her generous support of this research. HTV was funded by a CIHR Master’s Research Award, AM is funded by a CIHR Doctoral Research Award, and CWM was funded by a Vanier Canada Graduate Scholarship. Tissue banking in Montreal was supported by the Banque de tissus et données of the Réseau de recherche sur le cancer (RCC) of the Fond de recherche du Québec – Santé (FRQS), associated with the Canadian Tumor Repository Network (CTRNet). Tissue banking in Ottawa was supported in part by Ovarian Cancer Canada.

## Competing Interests

The authors declare that they have no competing interests.

## Data and Materials Availability

All data needed to evaluate the conclusions in the paper are present in the paper and/or the Supplementary Materials. The raw RNA-seq data will be made publicly available on GEO when the manuscript is published.

## Author Contributions

Conceptualization: HTV, AM, CWM, BCV

Methodology: HTV, AM, ORP, CM, DAL

Investigation: HTV, AM, ORP, CWM, EY, EM, ME, DAL

Resources: ML, EJ, TJF, DT, AMMM, RD, BCV

Data Curation: HTV, AM, EY

Writing—original draft: HTV, AM, BCV

Writing—review & editing: HTV, AM, ORP, CWM, EY, EM, AMMM, DAL, BCV

Visualization: HTV, AM, ORP

Supervision: DAL, BCV

Project administration: BCV

Funding acquisition: BCV

## Notes

### Competing Interest Statement

The authors have declared no competing interest.

### Summary of Updates

Figures 5 and 7 were revised; Author list was updated; Supplemental files updated

