## Supplementary material for "*BRCA* mutation alters the stromal landscape in normal ovaries": Brca Manuscript_Supplemental Figures

**Supplemental Table 1.** Patient information of the human ovary cohort. Case numbers marked with an asterisk (*) indicate samples that were used for FibroNest analysis. [#]vus – variant of uncertain clinical significance.

| Group | Case No. | Age | Genetic Mutation | Family History |
|---|---|---|---|---|
| **Premenopausal WT** | 1 | 44 | | |
| | 2 | 36 | | |
| | 3 | 40 | | |
| | 4 | 49 | | |
| | 5 | 47 | | |
| | 6 | 47 | | |
| | 7* | 41 | | Breast Cancer |
| | 8* | 45 | | |
| | 9 | 41 | | |
| | 10 | 43 | | |
| | 11 | 48 | | |
| | 12 | 44 | | |
| | 13* | 47 | | |
| | 14 | 49 | | |
| | 15 | 38 | | |
| | 16* | 49 | | |
| | 17 | 47 | | |
| | 18 | 46 | | |
| | 19 | 43 | | |
| | 20* | 49 | | |
| | 21 | 48 | | |
| | 22 | 49 | | |
| | 23 | 45 | | |
| | 24 | 47 | | |
| **Premenopausal BRCA** | 25* | 47 | BRCA1+ | Breast and Ovarian Cancer |
| | 26 | 36 | BRCA1+ | Breast and Ovarian Cancer |
| | 27 | 53 | BRCA2+ | Breast Cancer |
| | 28 | 49 | BRCA2+ | Breast Cancer |
| | 29 | 35 | BRCA1+ | Breast Cancer |
| | 30 | 37 | BRCA2+ | Breast Cancer |
| | 31 | 37 | BRCA2+ | Breast Cancer |

| | | | | |
|---|---|---|---|---|
| | 32* | 45 | BRCA2+ | Breast and Ovarian Cancer |
| | 33 | 38 | BRCA1+ | Breast and Ovarian Cancer |
| | 34 | 41 | BRCA2+ | Breast and Ovarian Cancer |
| | 35 | 39 | BRCA1+ | Breast and Ovarian Cancer |
| | 36 | 52 | BRCA1+ | Breast Cancer |
| | 37* | 45 | BRCA2+ | Pancreas and Colon Cancer |
| | 38 | 45 | BRCA1+ | Breast and Ovarian Cancer |
| | 39* | 35 | BRCA2+ | Breast Cancer |
| | 40 | 34 | BRCA2+ | Breast Cancer |
| | 41 | 43 | BRCA1+ | Breast Cancer |
| | 42 | 43 | BRCA2+ | Breast Cancer |
| | 43 | 39 | BRCA1+ | Ovarian Cancer |
| | 44* | 45 | BRCA2+ | Breast and Ovarian Cancer |
| | 45 | 42 | BRCA2+ | Breast and Ovarian Cancer |
| | 46 | 39 | BRCA1+ | Breast and Ovarian Cancer |
| | 47 | 42 | BRCA2+ | Breast and Ovarian Cancer |
| | 48 | 40 | BRCA2+ | Breast and Ovarian Cancer |
| **Postmenopausal WT** | 49* | 56 | | |
| | 50* | 62 | | |
| | 51* | 49 | | Ovarian and pancreatic cancer |
| | 52* | 58 | PMS2vus[#] | |
| | 53* | 77 | | Uterine and Colon Cancer |

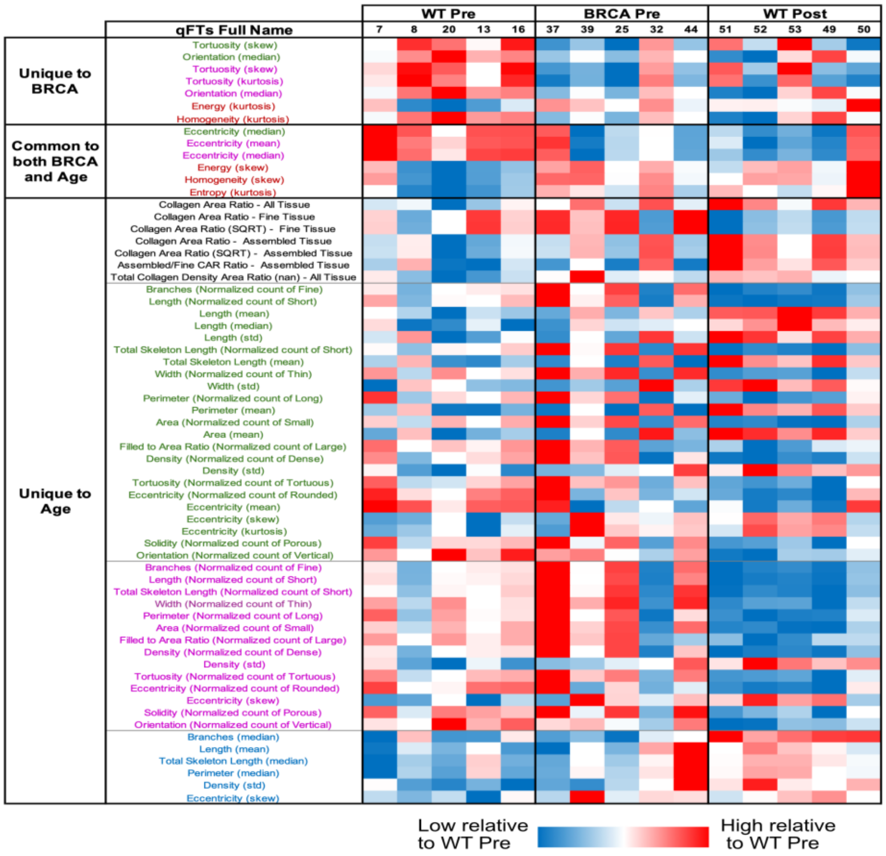


**Supplementary Fig. 1: Single fiber analysis using the FibroNest digital pathology platform identifies differences in ovarian cortex stratified by age and *BRCA* mutation.** Heatmap of fibrosis traits for each sample as identified by the FibroNest platform indicating directionality. Only significant quantitative fibrosis traits (qFTs; $p < 0.05$) are presented and are segregated based on being unique to BRCA, common to both BRCA and age, and unique to age. Numbers within each group identify patients as listed in Supplemental Table 1. Red indicates an increase in trait value score while blue indicates a decrease in trait value score amongst all groups. qFT colour descriptions are as follows: black – collagen content; green – collagen morphometry among all fibers; pink – collagen morphometry among fine fibers; blue – collagen morphometry among assembled fibers; red – fibrosis architecture. Significance was determined by Welch's t test (*n* =5).

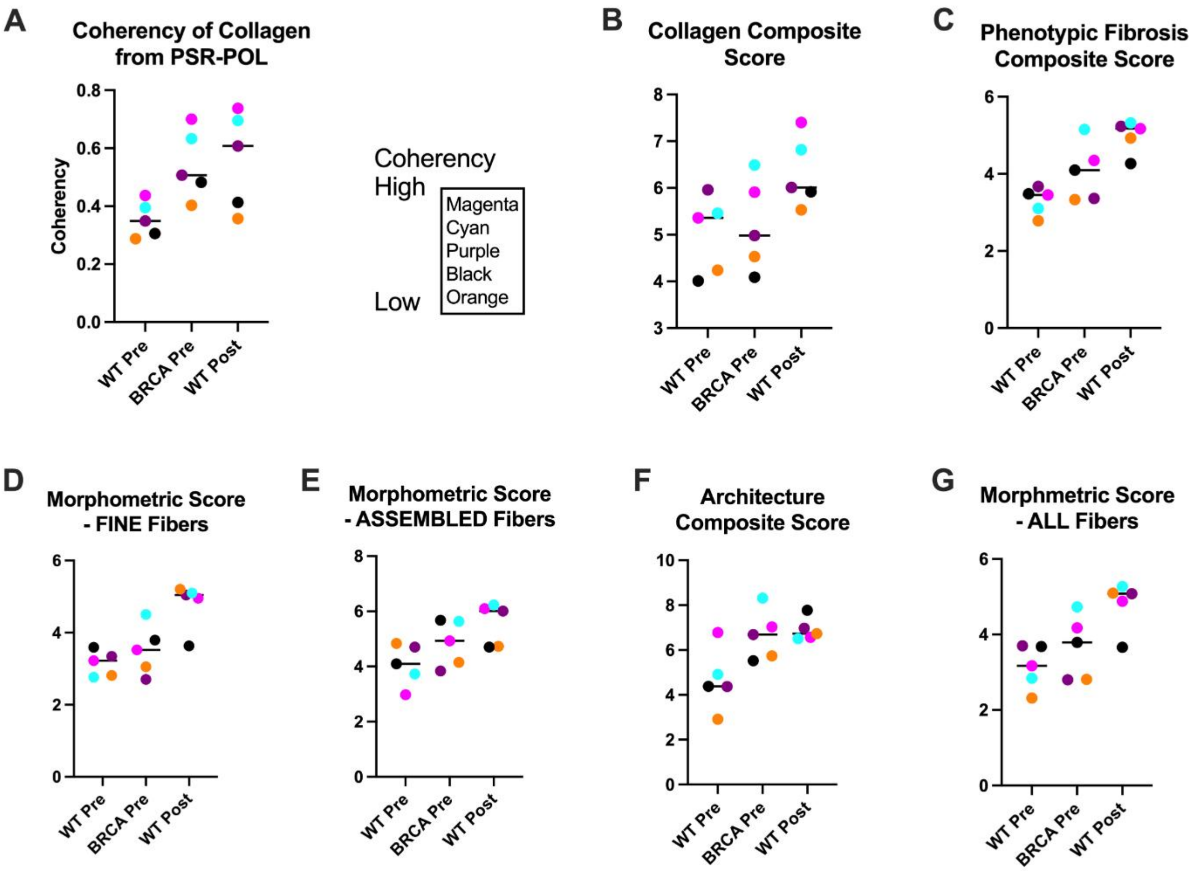


**Supplementary Fig. 2: Scores from FibroNest analysis show concordance with coherency values calculated by PSR-POL.** (A) Coherency values from PSR-POL for samples that underwent FibroNest analysis and ranked from high to low by colour (magenta – cyan – purple – black – orange). (B – F) FibroNest fibrosis scores by sample with PSR-POL value indicated by colour (*n* = 5).

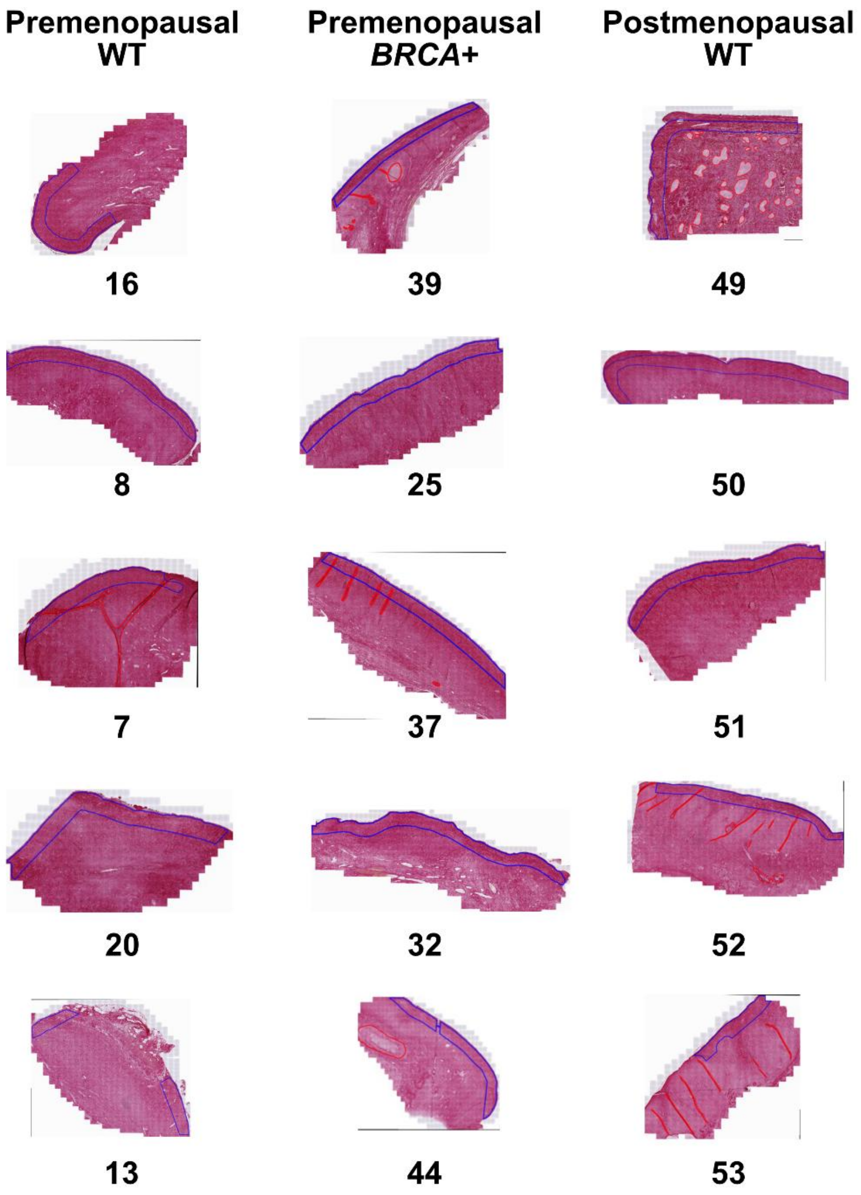


**Supplementary Fig. 3: Brightfield images taken of the ovarian cortex and submitted to the FibroNest platform for analysis.** Images were taken at 40X and then an ROI was designated for each sample beginning 30 μm below the surface and extending 300 μm in depth. Each sample is indicated with the patient case number that was identified in Supplemental Table 1.
